# Co-Stimulation–Induced AP-1 Activity is Required for Chromatin Opening During T Cell Activation

**DOI:** 10.1101/647388

**Authors:** Masashi Yukawa, Sajjeev Jagannathan, Andrey V. Kartashov, Xiaoting Chen, Matthew T. Weirauch, Artem Barski

## Abstract

Activation of T cells is dependent on organized and timely opening and closing of chromatin. Herein, we identify AP-1 as the transcription factor that directs most of this remodeling. Chromatin accessibility profiling showed quick opening of closed chromatin in naïve T cells within 5 hours of activation. These newly open regions were strongly enriched for the AP-1 motif, and indeed, ChIP-seq demonstrated AP-1 binding at more than 70% of them. Broad inhibition of AP-1 activity prevented chromatin opening at AP-1 sites and reduced expression of nearby genes. Similarly, induction of anergy in the absence of co-stimulation during activation, was associated with reduced induction of AP-1 and a failure of proper chromatin remodeling. The translational relevance of these findings was highlighted by the substantial overlap of AP-1–dependent elements with risk loci for multiple immune diseases, including multiple sclerosis, inflammatory bowel disease and allergic disease. Our findings define AP-1 as the key link between T cell activation and chromatin remodeling.

## Introduction

Upon encountering an antigen, naïve T helper cells are activated and differentiate over several days into various effector lineages that contribute to immune responses(O’Shea and Paul, 2010; Russ et al., 2013). These differentiated effector cells secrete different sets of cytokines and have specific functions in orchestrating immune responses against pathogens. In the contraction phase of the response, most effector cells die, but a few survive and become long-lived memory cells (Ben Youngblood et al., 2017). We and others have demonstrated that epigenetic states induced during T cell activation, differentiation, and memory formation are associated with T-cell lineage stability and plasticity, cytokine production in effector cells, and rapid recall response in the memory cells (Vahedi et al., 2012; Barski et al., 2009; Komori et al., 2015; Smith et al., 2009; Hawkins et al., 2013; Mukasa et al., 2010; Mazzoni et al., 2015; Sekimata et al., 2009; Wei et al., 2009; Ohkura et al., 2012). An outstanding question in the field is how the epigenetic changes are induced and targeted to specific loci during primary activation of T cells.

The differentiation of T cells is a multi-step process starting with T cell activation. The activation is accomplished through simultaneous stimulation of the T cell receptor (TCR) and co-stimulatory receptors such as CD28. Downstream NFATs, AP-1 (a heterodimer of FOS and JUN proteins), and NFκB are activated via Ca^2+^-calcineurin, MAP kinase and PI3K/PKC pathways (Fathman and Lineberry, 2007; Crabtree and Olson, 2002; Zhu and Paul, 2010; Jain et al., 1994; Rochman et al., 2015). Concurrently with activation signals, differentiation signals provided by the cytokine milieu lead to activation of JAK-STAT pathways, induction of lineage-specific transcription factors (TFs), and eventually lineage-specific cytokine gene expression (Zhu et al., 2010). The *Il2* locus has previously been used as a model to study activation-induced transcriptional regulation. The *Il2* promoter has several AP-1 and NFAT binding sites that are conserved between human and mouse (Rooney et al., 1995; Macian et al., 2001). The binding sites are adjacent, and AP-1 and NFAT form a heteromer (Jain et al., 1994; Chen et al., 1998) and synergize to induce *Il2* expression (Walters et al., 2013; Nguyen et al., 2010). Mutation of these binding sites prevents Il2 expression (Walters et al., 2013). NFκB and several other TFs also participate in *Il2* regulation during T cell activation via their binding sites near the promoter (Thaker et al., 2015; Skerka et al., 1995). However, the mechanisms of transcriptional regulation during T cell activation are not common for all genes. For example, *IL2* expression is dependent on new protein synthesis, but *IL10*, *IFNG*, and *TNF* are not (Sareneva et al., 1998).

Herein, we profiled chromatin accessibility during the early stages of T cell activation in human primary naïve CD4 T cells. We were struck by the massive number of regions undergoing remodeling within 5 h of activation and by the considerable enrichment of AP-1 motifs. Chromatin immunoprecipitation (ChIP-seq) demonstrated AP-1 binding at the majority of these regions, often together with its partner NFAT1. AP-1 was also strongly present at super enhancer (SE) elements formed during activation. Whereas prior studies have focused on genetic disruption of individual AP-1 members, herein we broadly blocked the AP-1 family in human naïve T cells by electroporating a dominant-negative protein (A-FOS); this resulted in loss of chromatin remodeling and T cell activation. Conversely, AP-1– associated chromatin changes were absent during induction of T cell anergy. The translational significance of these findings to clinical medicine was supported by the overlap of activation-specific enhancers and AP-1 binding sites with single-nucleotide polymorphisms (SNPs) associated with increased risk for a variety of diseases, most substantially found for multiple sclerosis.

## Results

### Characterizing open chromatin regions

Human naïve CD4 T cells isolated from the blood of healthy donors were activated with anti-CD3/CD28 beads for 5, 24, and 60 h. (Fig. S1 A and B). Open chromatin in resting and activated cells was profiled by ATAC-seq, and differentially accessible regions were identified (Fig. 1 A and B). Herein, we will refer to loci that are accessible only in naïve cells or only activated cells as “naïve open region (NOR)” or “effector open region (EOR)”, respectively, and to those loci maintained in both naïve and activated cells as “common open region (COR)”. For example, *KLF2*, encoding a quiescence-related TF (Kuo et al., 1997) downregulated upon activation, had a NOR, whereas *EGR2*, an early growth response TF that is induced during activation, has both a COR and an EOR (Fig. 1 A and B). Overall, genome-wide analysis indicates that the chromatin remodeling that occurs at the early stages of T cell activation is dramatic (Fig. 1 C): in addition to the 11,117 open regions present in resting T cells (NORs plus CORs), 10,218 regions became accessible after 5 h of activation (EORs). Only 899 of open regions closed. We observed fewer changes at later time points, indicating that chromatin remodeling is more active immediately after activation. For the 5-h time point, we also identified a high confidence set of regions that were consistently identified as NOR, COR, or EOR in T cell from two separate donors and were used for further analysis (Fig. 1 C). As expected, chromatin opening was associated with gene expression changes: genes nearest to EORs were enriched among genes upregulated during T cell activation (FDR = 0.018 for 736 genes with 1 EOR and FDR < 0.001 for 338 genes with > 1 EOR), (Fig. 1 D). In contrast to EOR genes, genes possessing NORs were downregulated during T cell activation (FDR < 0.001 for 95 genes with 1 NOR), (Fig. S1 D). Furthermore, we found that the set of genes in the vicinity of 5-h EORs (EOR genes) were enriched for genes related to T cell function and activation (GO terms, “lymphocyte activation” and “T cell activation”) by gene ontology analysis, whereas those genes that became accessible at later time points (5-24 h, 24-60 h) were involved in cell migration, proliferation, and metabolism (Fig. 1 E). We also found that NOR and COR gene sets were not enriched with genes related to T cell function and activation (Fig. S1 D). Further, we intersected the EORs with the genetic variants that related to various human conditions using the regulatory element locus intersection (RELI) approach (Harley et al., 2018). The results show that the risk SNPs for autoimmune or allergic diseases (e.g., Multiple sclerosis, Inflammatory bowel disease, Crohn disease, Self-reported allergy, and combined allergic disease SNPs (asthma, hay fever or eczema) have significant overlap with EORs (Fig. 1 F and Table S1). Thus, formation of open chromatin at early time points during T cell activation is likely to play an important role in T cell activation and its disruption may play a key role in immune disease processes.

**Figure 1.**
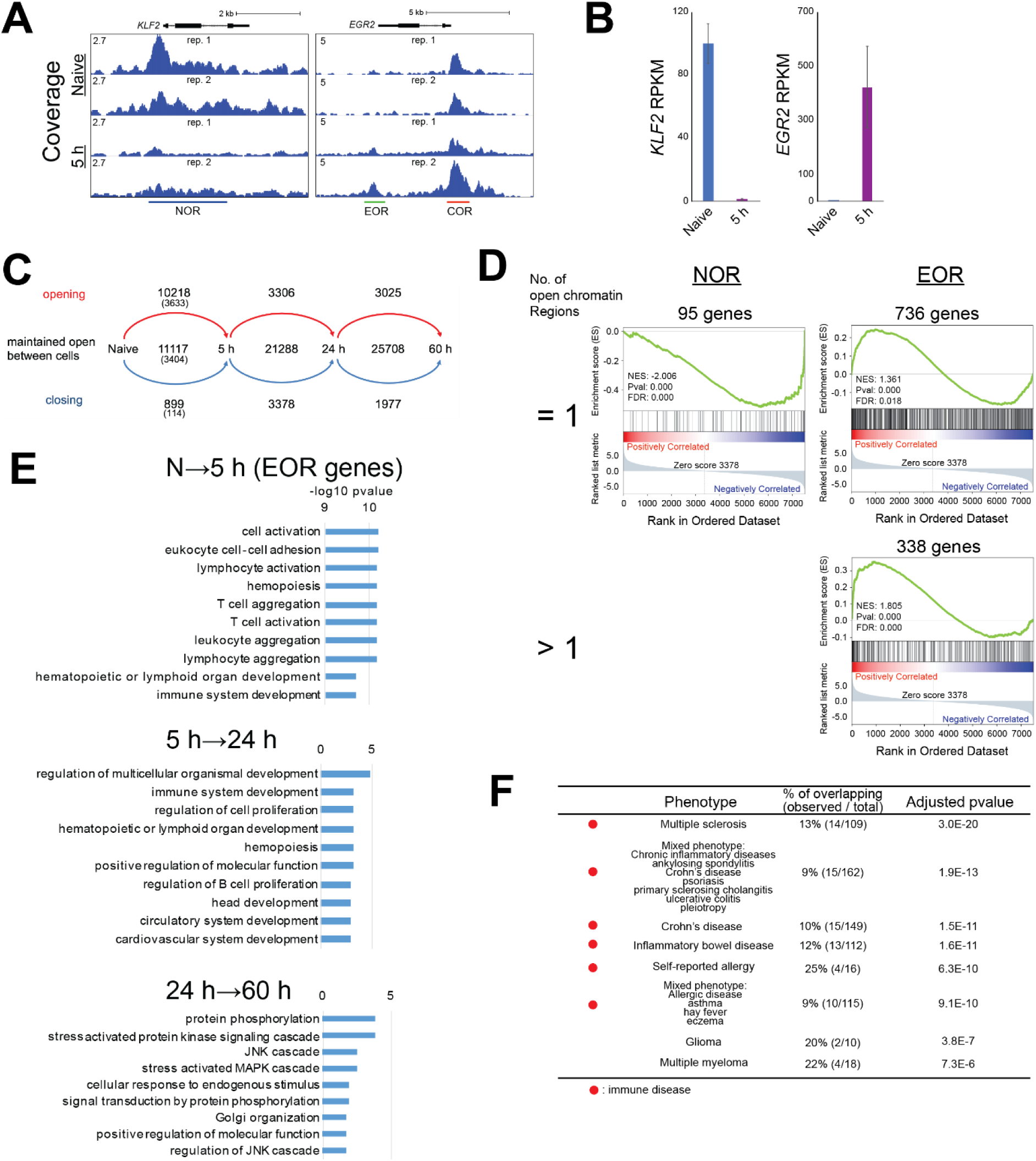
Characterization of open chromatin during T cell activation. (A) UCSC genome browser screenshots show open chromatin at the *KLF2* and *EGR2* loci in naive and activated T cells at 5 h. The y axis shows ATAC-seq coverage by estimated fragments normalized to the number of mapped reads. (B) Bar plots show expression of *KLF2* and *EGR2* genes by RNA-seq. Mean and standard error are shown. n=2. (C) Flow diagram shows the dynamics of open chromatin regions identified in naive T cells and T cells activated for the indicated time period. The numbers indicate the number of newly opened, maintained open, or newly closed chromatin regions. Numbers in parentheses indicate the number of highly reliable regions used for further analyses. (D) Gene set enrichment analysis (GSEA) compares the gene list ranked by expression fold change during activation with the sets of genes that are located next to 1 or more than 1 NOR or EOR. NES: Normalized Enrichment Score. (E) Gene ontology analysis of genes adjacent to chromatin regions that open during T cell activation. Top gene ontology (GO) biological processes terms and −log10 p-values are shown. N: naïve. (F) Overlap between disease risk SNPs and EORs. Significance of overlap between disease risk SNPs and EORs as calculated by RELI approach. Only top 10 GWAS term is shown. Full list for NOR, COR, and EOR is available in Table S1.

Interestingly, the majority of NOR and COR loci were found in promoter regions, whereas the majority of EOR loci were not (Fig. 2 A). These findings suggest that these newly accessible sites might serve as regulatory elements. To investigate their function, we examined changes of histone modifications at these open chromatin regions (Fig. 2 B and C). For example, at the *CD82* gene, we observed increased accessibility at two internal elements upon activation, and this was accompanied by an increase in H3K27Ac levels, suggesting that the elements may function as enhancers. Genome-wide analysis showed that the levels of H3K27Ac and H3K4me3 at the EOR were significantly increased in the transition from naïve to activated cells at 5 h (Fig. 2 C). These two histone modifications are considered to be hallmarks of enhancers and strong enhancers, respectively (Ernst et al., 2011). In conjunction with the EOR association with transcriptional upregulation, these findings suggest that the EOR are likely to serve as transcriptional enhancers.

**Figure 2.**
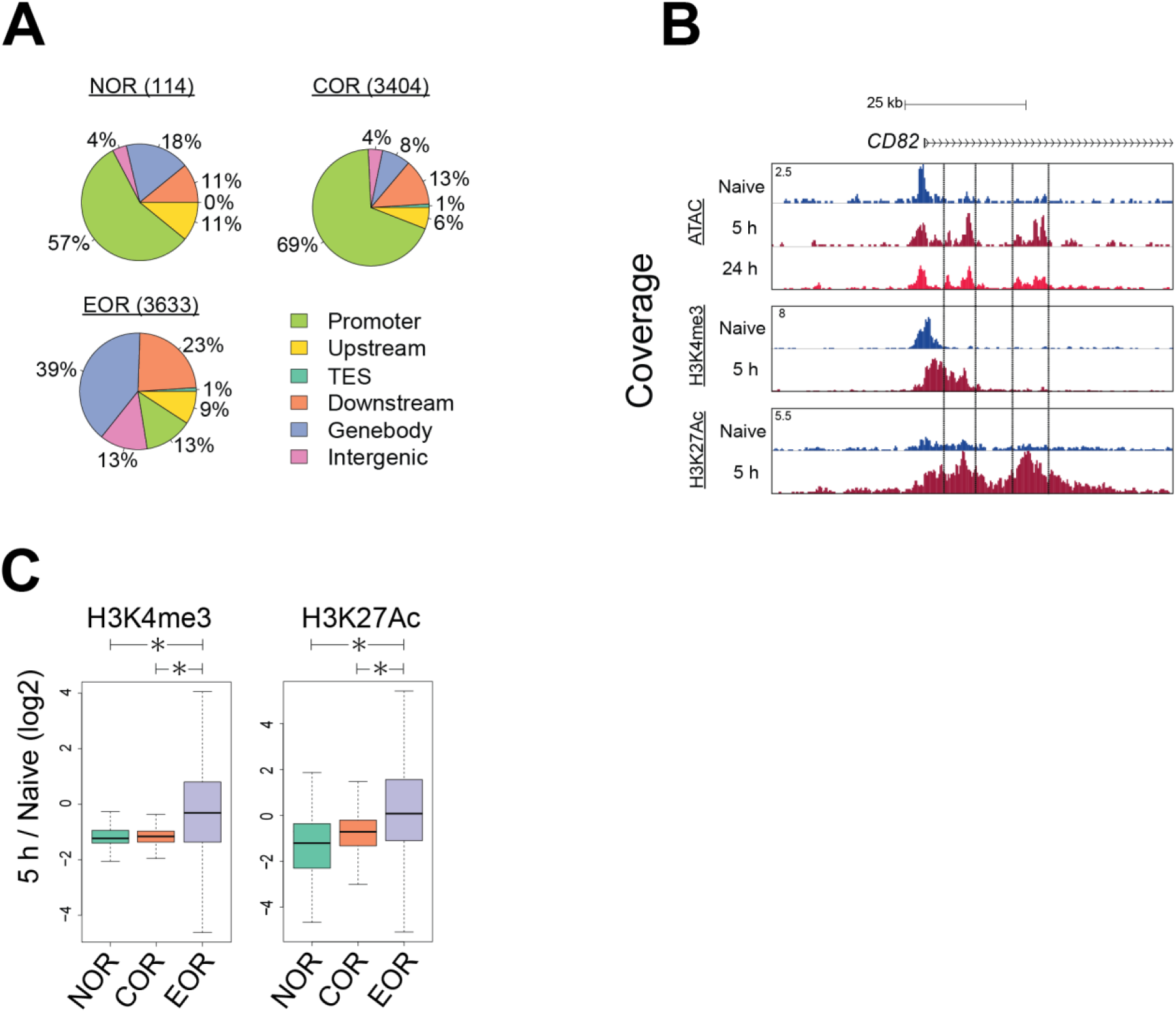
EORs function as transcriptional enhancers during T cell activation. (A) Pie charts show the number and distribution of open chromatin areas relative to gene locations. (B) UCSC genome browser screenshot of ATAC-seq and ChIP-seq for H3K4me3 and H3K27Ac at the *CD82* locus. The y axis shows the ATAC/ChIP-seq coverage by estimated fragments normalized to the number of mapped reads. (C) Changes in the H3K4me3 and H3K27Ac levels at the open chromatin regions during T cell activation. The y axis shows the log_2_ of the normalized ratio of ChIP-seq signals between naive and 5 h activated T cells (5 h/Naive). *p < 0.01 (Wilcoxon rank-sum test).

### Transcription factors AP-1 and NFAT1 bind effector open chromatin regions

To identify TFs that may play a role in chromatin remodeling during activation, we applied the HOMER software (Heinz et al., 2010) to identify TF binding sites over-represented within the differentially accessible regions (Fig. 3 A and Fig. 2 A and B). Analysis showed that DNA motifs of AP-1 or the AP-1/NFAT composite element were enriched in EOR but *not* in NOR and COR. Interestingly, the NFAT motif alone was not enriched as strongly. BORIS and CTCF motifs were prominent at the EOR sites not bound by AP-1 (Fig. S2 C), suggesting that nuclear organization may be also affected by activation-induced chromatin opening. In contrast to EORs, NORs were enriched in motifs for EGR and KLF, which have been reported to be negative regulators of T cell activation (Safford et al., 2005) and to maintain the naïve state in T cells (Yamada et al., 2009). For COR, motifs of ETS, which is important in T cell development and activation (Panagoulias et al., 2016; Muthusamy et al., 1995), were prominent. Indeed, enrichment of JunB motifs and binding at the sites of chromatin remodeling was previously reported in mouse T cells (Bevington et al., 2016). To test whether AP-1 and NFAT TFs actually bind EORs in human T cells, we performed ChIP-seq for cFOS and JUNB (components of AP-1), NFAT1, NFAT2, NFκB, and cMyc, which are known to be important for T cell activation and proliferation (Liu et al., 2016; Trushin et al., 1999; Wong et al., 1999; Chou et al., 2014; Wang et al., 2011). cFOS, JUNB, and NFAT1 were present in EORs more than in NORs and CORs, whereas NFκB and cMyc were not enriched in EORs (Fig. 3 B). For example, the *TNFRSF18* gene locus had two EOR regions (Fig. 3 C). One of them was bound by AP-1 and NFAT1, but the other was only bound by AP-1. Previously, it was reported that AP-1 and NFAT formed a heteromer on the *IL2* promoter (Chen et al., 1998), leading us to examine combinatorial effects of the TFs on the opening of chromatin genome-wide (Fig. 3 D). Remarkably, more than 70% of EOR were bound by AP-1 (cFOS/JUNB) alone or AP-1 and NFAT1 together. Further, almost all NFAT1-bound EOR were also bound by AP-1 (cFOS or JUNB). Interestingly, the triple combination of cFOS, JUNB, and NFAT1 resulted in the highest accessibility, followed by the cFOS/JUNB dimer, whereas NFAT1 sites not bound by AP-1 showed minimal opening (Fig. 3 E) in agreement with the NFAT-only motif not being enriched in EORs. Moreover, the H3K27Ac level was also the highest in regions with the triple combination (Fig. S3 A). Furthermore, the level of chromatin openness was correlated with the strength of cFOS, JUNB, and NFAT1 binding, (Fig. S3 B). These data suggest either a role for AP-1 in chromatin remodeling during T cell activation or passive binding of AP-1 complexes to enhancers opened by other TFs.

**Figure 3.**
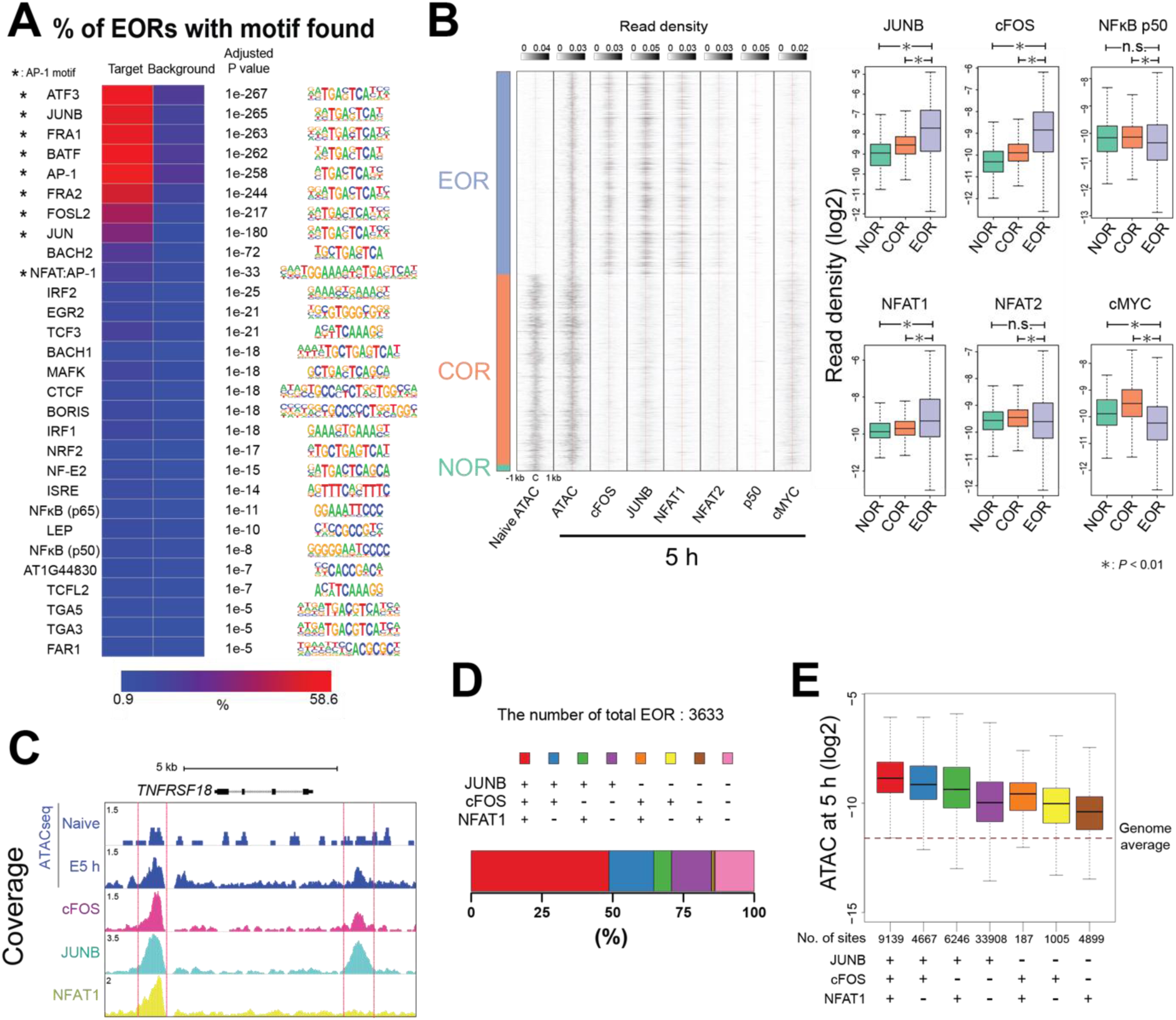
NFAT1 and AP-1 bind to EORs. (A) DNA binding motifs enriched in EORs. Heatmap shows the percentage of EORs with motifs. Overrepresented motifs were identified by HOMER analysis and selected with an adjusted p ≤ 10^−5^ and target / background > 2. *: AP-1 motifs (B) NFAT1 and AP-1 bind to EORs. Left: Fragment density heatmaps show read density of the ATAC-seq and TF ChIP-seq signal at open chromatin regions. C: Center of open chromatin regions. Right: Box plots show the TF ChIP-seq read density at open chromatin regions in activated T cells at 5 h. (C) UCSC genome browser screenshot showing ChIP-seq for AP-1 and NFAT1 at the *TNFRSF18/GITR* locus. Regions marked with red lines are EORs. (D) Stacked bar plot showing the percentage of EORs overlapping with significant TF ChIP-seq peaks. (E) Boxplot showing the ATAC-seq signal in activated T cells at 5 h in the regions bound by a given combination of TFs. The genome average indicates the average ATAC-seq signal for the whole genome. The y axis indicates ATAC-seq tag density over peaks.

### Formation of super enhancers involves chromatin opening

Combinations of several nearby enhancer elements with unusually high total level of H3K27 acetylation and/or binding of Mediator or BRD4 proteins are known as super enhancers (SEs) and are believed to induce expression of cell type–specific genes (Whyte et al., 2013; Selective Inhibition of Tumor Oncogenes by Disruption of Super-Enhancers, 2013). Therefore, we examined whether EORs were involved in the assembly of SEs. We identified 640 and 384 SEs in naïve and activated T cells, respectively, and 203 of them were shared between the cell states (Fig. S4 A and B). As expected, the presence of SEs resulted in high expression of nearby genes (Fig. S4 C). SE formation during T cell activation was associated with chromatin remodeling. For example, the *IL2RA* SE formation was accompanied by open chromatin formation after activation (Fig. 4 A and B). In addition to the EOR frequency being higher in activated SE (28.3 EOR/ 1 Mb), the frequency of EORs also was higher in shared SE (24.4 frequency EOR/ 1 Mb) (Fig. 4 C). SEs in naïve cells seem to rely on NOR and COR in contrast to SE in activated cells (Fig. 4 C and Fig. S4 D and E). Furthermore, AP-1 and NFAT1 were bound to activated and shared SEs at high frequency in contrast to naïve SEs (Fig. 4 D). Collectively, these results suggest that AP-1 and NFAT1 likely play a role in the formation of SEs during T cell activation.

**Figure 4.**
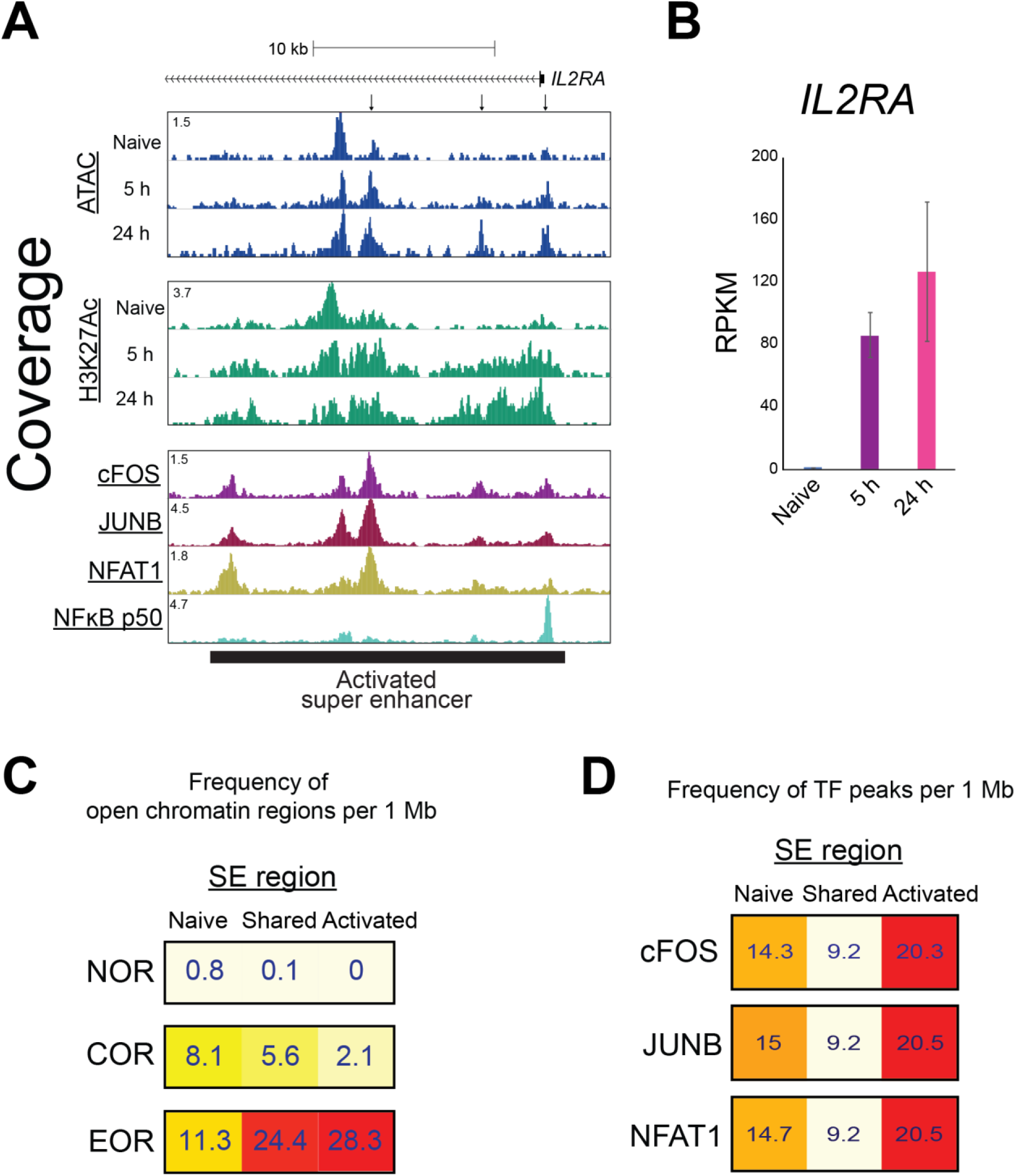
SE formation is associated with open chromatin formation. Super enhancers were identified using H3K27Ac ChIP-seq. (A) UCSC genome browser screenshot showing the ATAC and ChIP-seq signal for H3K27Ac and TFs around the super enhancer located in the *IL2RA* locus. (B) Bar plots show expression of *IL2RA* by RNA-seq. Mean and standard error are shown. n=2. (C) Heatmap showing the frequency of open chromatin regions per 1 Mb of a super enhancer. The frequency was calculated for super enhancers formed only in naive cells (Naive), only in activated cells (Activated), or in both cell types (Shared). (D) Heatmap showing the frequency of cFOS, JUNB, and NFAT1 peaks in super enhancer regions.

### AP-1 activity is required for open chromatin formation during T cell activation

In order to examine the role of AP-1 in the formation of open chromatin, we sought to inhibit AP-1 activity during T cell activation. Unfortunately, due to co-expression of multiple JUN and FOS family members in T cells and their common upregulation during activation (Fig. 5 A) (Jain et al., 1994), knock-out or knock-down strategies are unlikely to be successful. For this reason, we instead used the AP-1 dominant-negative protein A-FOS (Biddie et al., 2011). A-FOS can sequester JUN isoforms and prevent FOS-JUN complex formation and DNA binding. Bacterially expressed A-FOS or GFP control proteins were purified and electroporated into naïve T cells before activation. Electroporation efficiency was close to 100%, and GFP electroporation did not affect T cell activation (data not shown). We observed that in the A-FOS–electroporated cells, chromatin opening was decreased at EORs but not at NORs and CORs (Fig. 5 B). This blocking effect was more significant in EORs strongly bound by AP-1 (Fig. 5 C), and the effect appeared to be site-specific: an EOR bound by AP-1 was affected in the *IRF8* locus, whereas an EOR not bound by AP-1 was not affected (Fig. 5 D). The specific effect of A-FOS protein transduction provides reassurance that the electroporation procedure was functional and well received by the T cells. Furthermore, inhibiting AP-1 resulted in downregulation of activation-inducible genes such as *TBX21*, *IFNG*, and *CSF2* (Fig. 5 E). These results indicate that AP-1 binding is required for opening chromatin during T cell activation at many genomic loci.

**Figure 5.**
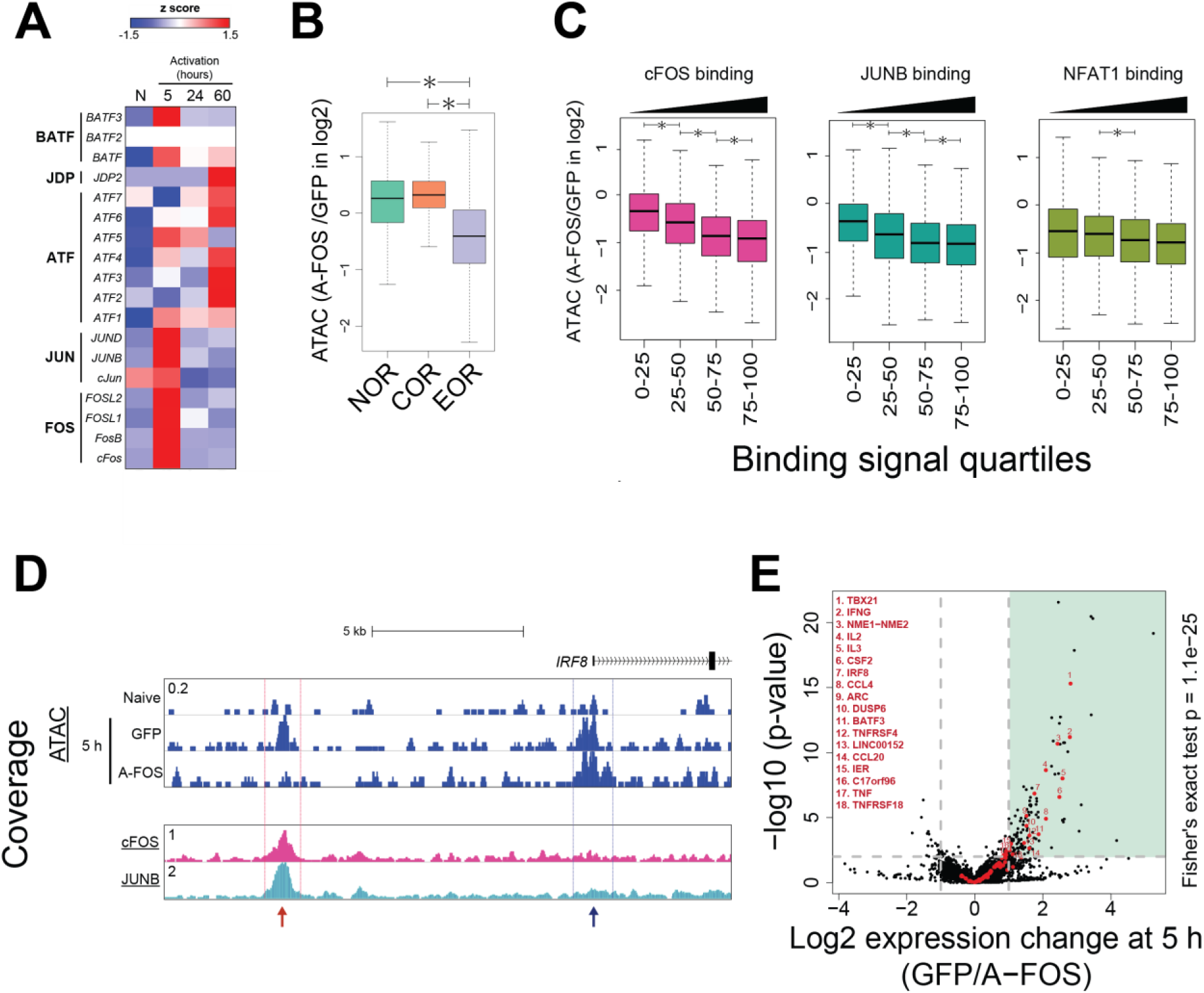
A-FOS, a dominant-negative regulator of AP-1, inhibits EOR formation. (A) Heatmap shows changes in mRNA expression level of AP-1 (FOS, JUN, ATF, and JDP) and BATF family proteins during T cell activation by RNA-seq. Color shows z-score. N: naive T cells. (B) Box plot showing the ratio of ATAC-seq signal in open chromatin regions between T cells electroporated with GFP and A-FOS after 5 h of activation. *p < 0.01 (Wilcoxon rank-sum test). (C) EORs were separated into quartiles on the basis of TFs’ ChIP-seq read density within peaks. Box plot shows the ratio of ATAC-seq signal in open chromatin regions between T cells electroporated with GFP and A-FOS after 5 h activation. *p < 0.01 (Wilcoxon rank-sum test). (D) UCSC genome browser screenshot showing the ATAC signal at the *IRF8* locus in naive cells and activated cells with GFP or A-FOS. The red arrow indicates an EOR dependent on AP-1. The blue arrow indicates an EOR independent of AP-1. (E) Volcano plot showing the changes in gene expression between GFP- and A-FOS– electroporated T cells by RNA-seq. X axis: ratio of expression between GFP and A-FOS T cells (log2). Y axis: −log10 DEseq2 p-value for the differential expression. Red dots: activated genes (Naïve→ 5-h Activation). Black dots: other genes. The vertical gray lines are at −1 and 1 on the x axis. The horizontal gray line is at 2 on the y axis. The p-value for overlap between genes induced during T cell activation and suppressed by A-FOS electroporation is calculated by the Fisher’s exact test.

### Co-stimulation is required for open chromatin formation

In order to better understand the role of AP-1–induced chromatin remodeling in the greater context of the immune response, we next performed T cell activation in the absence of CD28 co-stimulation (Fig. 6 A). Previous studies have shown that the lack of co-stimulation reduces activation of AP-1 and eventually leads to the induction of anergy (Macian et al., 2002; Kriegel et al., 2009; Rochman et al., 2015). Indeed, cells not receiving CD28 co-stimulation (Anergy) showed dramatic reduction of nuclear translocation of AP-1 and to a smaller degree NFκB p50, whereas nuclear levels of NFATs and NFκB p65 were reduced only slightly (Fig. S6 A). Genome-wide, the reduction of AP-1 activity was accompanied by decreased chromatin opening at EORs (Fig. 6 B and C). For example, in the absence of CD28 signaling, the AP-1– bound EOR in the vicinity of the *JAK2* gene was not remodeled and did not become acetylated on H3K27 (Fig. 6 D); consequently, *JAK2* demonstrated reduced gene expression (Fig. 6 E). The promoter-associated COR, also bound by AP-1, remained but with decreased H3K27Ac levels. Interestingly, NFAT binding was still detectable at the majority of sites including those that lost the ATAC signal in the absence of co-stimulation, suggesting that NFAT is not sufficient for opening chromatin (Fig. 6 D and Fig. 5S B). Failure of EOR formation in co-stimulation– deficient cells was especially prominent in regions that have higher openness among EORs in effector-activated cells at 24 h (Fig. 6 C). We tested whether failure of chromatin remodeling due to the lack of CD28/AP-1 affected the histone modifications and transcriptome in cells that did not receive co-stimulation. Indeed, failure of open chromatin formation led to reduced H3K27Ac level and gene expression at nearby genes (5 h, Spearman R = 0.13, p = 4.4e-15; 24 h, Spearman R = 0.32, p = 3.6e-87) (Fig. S5 C and D). These results suggest that induction of anergy by activation in the absence of co-stimulation is associated with the failure of AP-1–dependent chromatin remodeling.

**Figure 6.**
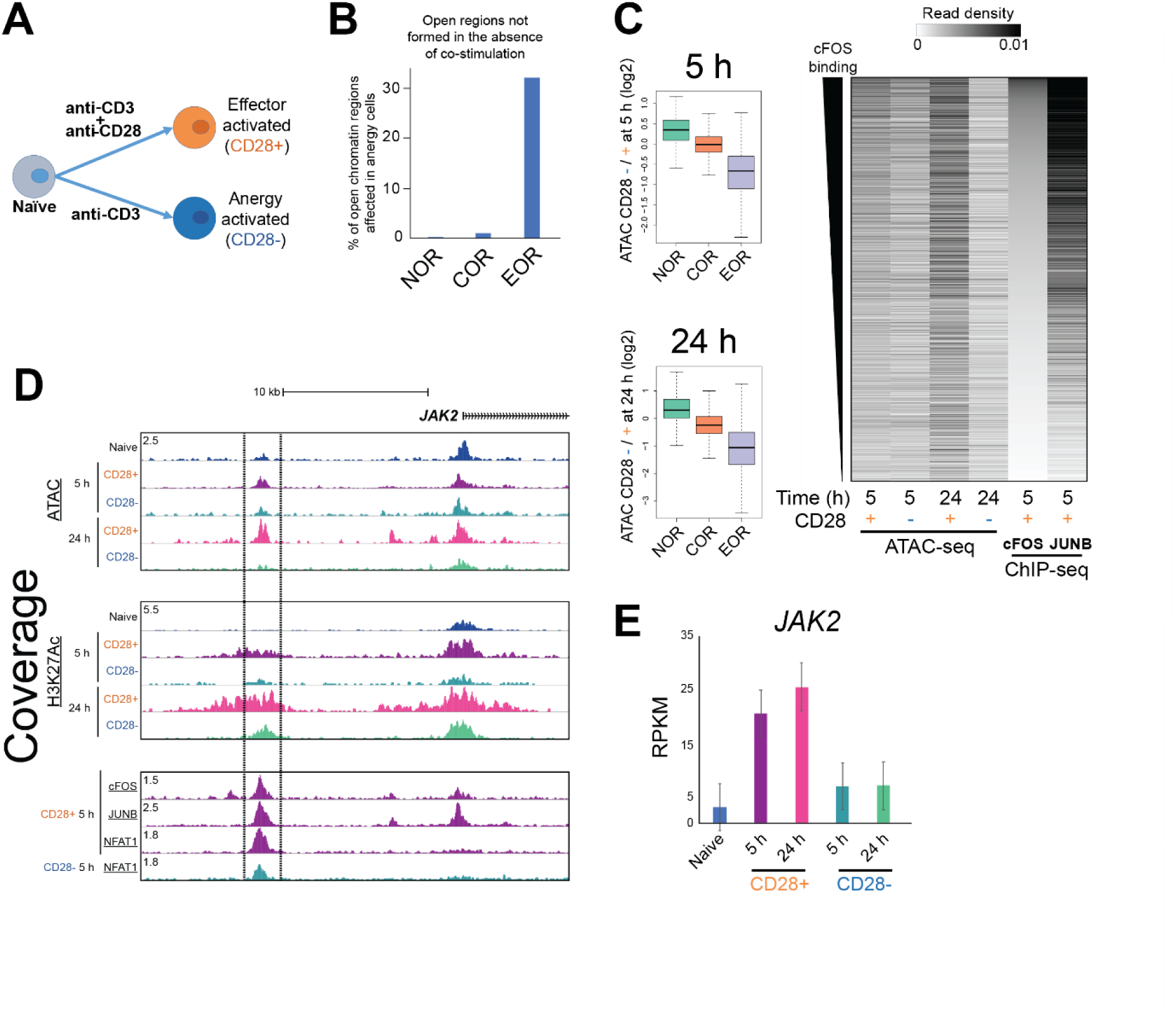
Incomplete open chromatin formation during induction of T cell anergy in the absence of co-stimulation. (A) Experimental approach: Naïve cells were activated with beads covered with anti-CD3 and anti-CD28 or only anti-CD3 antibodies. (B) Bar plot showing the percentage of open chromatin regions not formed at 5 h in cells that did not receive co-stimulation. (C) Chromatin fails to open in T cells that lack co-stimulation. Left: Box plot showing the ratio of ATAC signal between cells activated without and with CD28 co-stimulation at 5 and 24 h. Right: Heatmap showing ATAC-seq and TF ChIP-seq read density in EORs. EORs were sorted based on cFOS signal. (D) UCSC genome browser screenshot showing the ATAC-seq and ChIP-seq for H3K27Ac, AP-1, and NFAT1 in the *JAK2* locus. The region bordered by lines is open in fully activated cells but not in cells lacking co-stimulation. (E) Bar plot shows expression of *JAK2* by RNA-seq. Mean and standard error are shown. n=2.

## Discussion

T cell activation is associated with dramatic chromatin de-condensation that is essential for T cell differentiation and function (Rawlings et al., 2010; Lee et al., 2015). Genome-wide regulatory elements are bound by TFs that recruit chromatin-remodeling complexes and chromatin-modifying enzymes, leading to expression of activation-related genes. We mapped putative regulatory elements in human naïve and activated CD4 T cells by ATAC-seq and identified the regions that undergo remodeling at early time points during T cell activation. Many of them stay open in the resting memory cells. These regions also gained “positive” histone modification, such as H3K4me3 and H3K27ac, whereas the nearby genes tended to demonstrate increased expression. Interestingly, these regulatory elements showed significant overlap with SNPs associated with a number of immune diseases, underscoring their importance for the regulation of the immune system. Next, we identified the TF binding motifs that are present at the sites of remodeling and thus may control chromatin remodeling at these elements. To confirm the results of the computational analysis, we monitored actual binding of NFAT1 and AP-1, as well as NFκB and cMYC, during activation by ChIP-seq. Comparing TF binding and chromatin opening data suggested that AP-1 is the factor responsible for opening chromatin at regulatory elements during T cell activation. Indeed, delivery of the AP-1 dominant-negative protein A-FOS into naïve T cells resulted in the T cells’ failure to remodel chromatin. Finally, we found that lack of the AP-1 activation during induction of anergy resulted in a failure of T cells chromatin remodeling.

It has long been known that activation of T cells requires two signals. Work from the Rao laboratory and others has shown that insufficient induction of AP-1 in the absence of co-stimulation fails to induce *Il2* gene expression and eventually leads to anergy (Macian et al., 2002; Kriegel et al., 2009; Rochman et al., 2015). However, murine cFOS knock-out T cells demonstrated normal response to activation, suggesting that either cFOS was not necessary (Jain et al., 1994) or that the other FOS family members, such as FOSB, FRA-1, and FRA-2, could substitute for cFOS (Fleischmann et al., 2000; Gruda et al., 1996). Indeed, all AP-1 family genes are dramatically upregulated upon T cell activation (data not shown). Given that the AP-1 family includes 18 genes, generating a complete deletion was impractical. Furthermore, manipulating expression in resting naïve T cells is difficult because successful viral transduction requires T cell proliferation, and although plasmid electroporation is possible, the process impairs cell activation (data not shown). We overcame this dual problem by (i) employing the A-FOS dominant-negative protein that sequesters JUN family members into unproductive A-FOS/JUN complexes (Biddie et al., 2011) and (ii) electroporating the *protein* into resting naïve T cells, which (unlike plasmid electroporation) did not affect T cell activation. In the presence of A-FOS, T cells failed to remodel their chromatin at multiple AP-1–bound sites. This experiment explains the role of AP-1 in T cell activation—establishing chromatin remodeling. Without CD28-driven AP-1 induction, naïve CD4 T cells could not establish open chromatin regions to act as enhancer elements for inducing gene expression in T cell activation. Our results further confirmed the hypothesis (Jain et al., 1994) that although individual FOS family proteins are redundant, induction of the AP-1 family is critical for T cell activation.

CD4 T cells control functions of other immune cells and are the key regulators of immune response and homeostasis. Impairment of T cell activation may lead to persistent infections and cancer, and over activation of T cells may cause autoimmune or atopic disease. Aside from elucidating the epigenetic mechanism of T cell activation, our data demonstrate a significant overlap between putative regulatory elements employed in T cell activation (EORs) with previously unexplained risk loci for several autoimmune diseases, such as Crohn disease, multiple sclerosis, and inflammatory bowel disease (Fig. 1 F) (Soderquest et al., 2017). Previously AP-1 sites were found to be enriched near SNPs for autoimmune diseases (Farh et al., 2014). In order to gain further clues into potential disease mechanisms, we repeated the RELI analysis with TF ChIP-Seq data in order to identify the candidate TFs that may bind risk loci. This analysis revealed that risk loci for multiple immune diseases were enriched for AP-1– and NFAT1 (Fig. 7 A-C). We next endeavored to check whether the EORs overlapping disease risk loci are differentially open (or modified) in T cell from patients. Although we could not find such data for our top hits (MS and IBD), a recent study (Seumois et al., 2014) reported H3K4me2 profiles for Th2 cells obtained from asthmatics and healthy controls. We found that 2 allergic disease risk loci (CD25 and PHF19) that significantly overlap with EORs (Fig. 7C) showed significant increase in the level of H3K4me2 in asthma patients compared to healthy controls (FDR<0.1, Fig. 7 D). These findings indicate the importance of proper chromatin remodeling for immune homeostasis.

**Figure 7.**
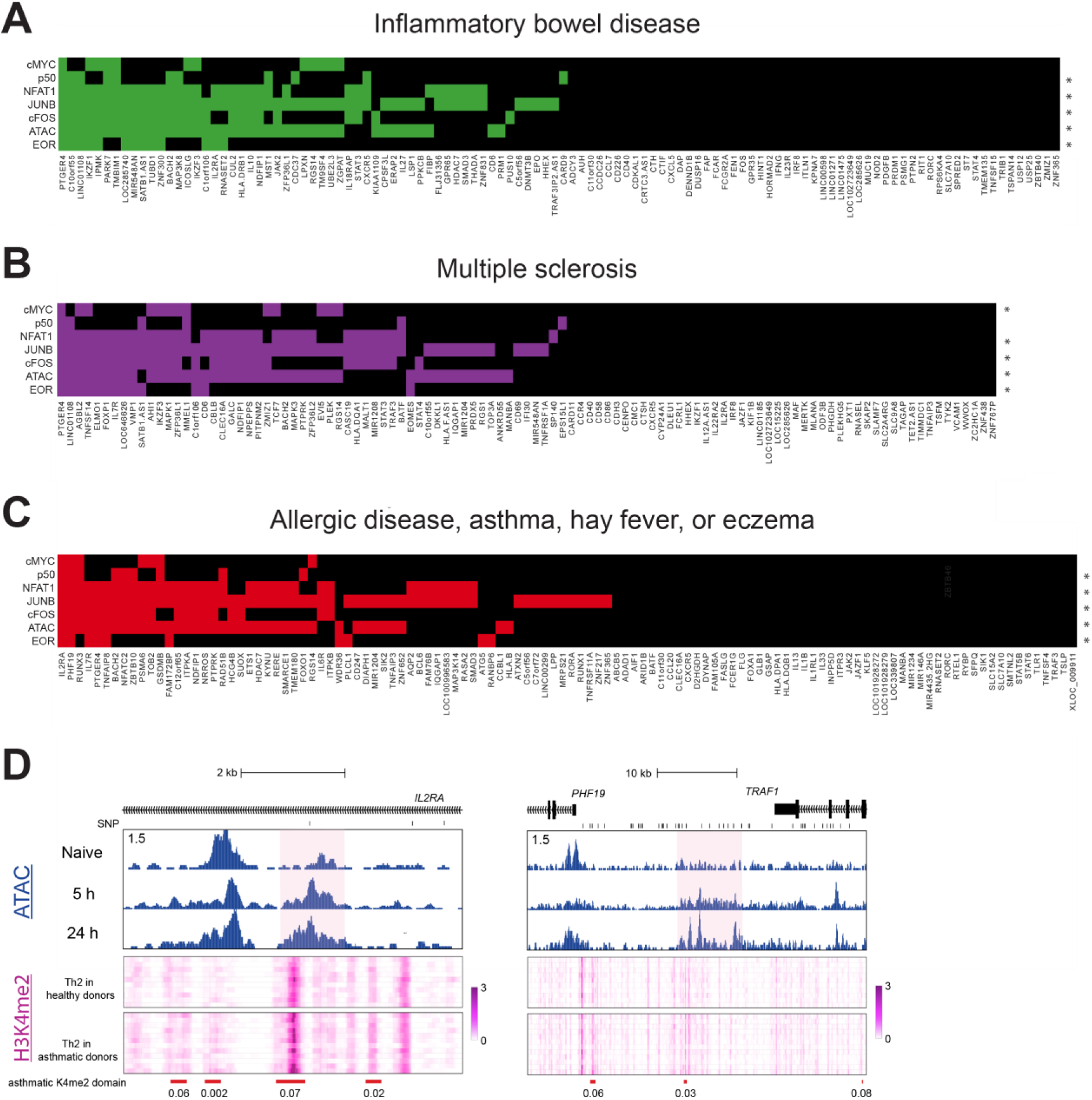
Intersection of immune disease risk loci, open chromatin regions and TF binding interactions with the genome. (A-C) Heatmaps for IBD (A), MS (B), and Allergic disease, asthma, hay fever, or eczema (C) risk loci. The x axis displays diseases-associated loci. An asterisk means the genomic coordinates of the ChIP-seq peaks measuring open chromatin regions or TF binding (at 5 h) significantly intersect with disease risk loci at a RELI corrected p-value threshold of 10^−6^. A colored box indicates that the given locus contains at least one disease-associated variant located within an open chromatin region or a ChIP-seq peak for the given TF. (D) Genome browser screenshot showing the ATAC-seq in naïve and activated T cells and heatmap showing H3K4me2 level in Th2 cells of asthmatics and healthy controls in the *IL2RA* and *PHF19* loci. Red bars indicate differentially enriched loci (FDR is shown under the bars). Pink backgrounds indicates EORs.

How does binding of AP-1 lead to chromatin remodeling? Previous studies have shown that AP-1 proteins can recruit components of BAF chromatin remodeling complex. In particular, SMARCD1 was shown to be AP-1 interacting partner in yeast two-hybrid screen(Ito et al., 2001). ATF3 was shown to physically interact with SMARCA4 (BRG1-ATPase) (Xu et al., 2011), while a more recent study showed interaction of FOS with 10 out of 15 BAF members and ability of other FOS family members (FOSD, FOSL1/2) to recruit both SMARCB1 and SMARCD1 (Vierbuchen et al., 2017).

Other TFs are involved in regulating the epigenome during T cell activation. Recent studies discussed the roles of STATs, GATA3, and T-BET (Vahedi et al., 2012; Durant et al., 2010; Wan et al., 2015; Pham et al., 2013; Kanhere et al., 1AD). Moreover, although AP-1 is required for chromatin remodeling at the regulatory elements for T cell activation, our study does not completely answer the question of specificity: many cell types express AP-1 and NFATs but do not induce T cell genes such as *IL2* (Armstrong, 2004; Turner et al., 1998; Ranger et al., 2000; Jauliac et al., 2002). Thus, the specificity would need to be established either by a pre-existing epigenetic state or by interaction with additional TFs. For this reason, AP-1 cannot be considered to be a pioneer factor (Zaret and Mango, 2016) just by itself. RUNX family proteins, which are broadly expressed in lymphocytes and regulate expression of *IL2* (Djuretic et al., 2006; Ono et al., 2007) and ETS family members (Bevington et al., 2016) may play a role, but involvement of additional TFs is also possible.

## Method

### Isolating, culturing, and activating human CD4 T cells

Blood filters containing cells from de-identified donors were supplied by Hoxworth Blood Center at the University of Cincinnati. Peripheral blood mononuclear cells (PBMCs) were isolated by Ficoll density gradient centrifugation. Isolated PBMCs were processed with the EasySep™ Human Naïve CD4^+^ T Cell Isolation Kit (STEMCELL, cat#19555) to negatively isolate human CD4^+^/CD45RO^−^ cells (naïve T cells), and the purity was confirmed by flow cytometry (Fig. S1 B). Isolated naïve T cells were cultured in RPMI media with L-glutamine (HyClone, cat#SH30027.01) supplemented with 10% FBS and 25 μM 2-ME before activation. For bead activation, 100 μL of anti-mouse pluriBeads (pluriSelect, cat#31-GTaMS-10) were incubated with 24 ng of anti-human CD3 antibody (Bio X Cell, cat#BE0001-2) and 52 ng of either anti-human CD28 antibody (Bio X Cell, cat# BE0248) or mouse IgG (Millipore, cat#12-371) with rotation for 3-4 h at RT. The beads were centrifuged and washed with PBS twice. Antibody-bound pluriBeads were directly added to the cultures of human CD4^+^ T cells, and the cultures were shaken for 4 hr and swirled every 10 min until cells were bound to the beads.

### Flow cytometry assay

Cells were stained with PE-Cy™7 Mouse Anti-Human CD45RO, FITC Mouse Anti-Human CD45RA, and FITC anti-human CD69 antibodies. Flow cytometry was performed on the BD FACS Canto II using the BD FACSDIVA.

### Isolating nuclear proteins and western blotting

T cells were collected and lysed with nuclear isolation buffer (10 mM HEPES-KOH [pH 8.0], 10 mM KCl, 1.5 mM MgCl_2_, 0.34M sucrose, 10% glycerol, 1 mM DTT, and 1X protease inhibitors) for 10 min with shaking every 3 min. The nuclear pellet was collected by centrifugation and was suspended and washed with nuclear isolation buffer twice. The nuclear pellet was directly lysed in SDS sample buffer with 8M urea. SDS-PAGE was carried out in the Bolt® Bis-Tris system (Thermo Fisher Scientific). Transfer was carried out with 120 V for 90 min to a PVDF membrane. Blocking of the membrane was performed with Odyssey Blocking Buffer (LI-COR, cat#927-50000) for 60 min at RT with shaking. The antibody incubation was carried out at 4 °C overnight with an orbital shaker. Antibodies to cFOS (#2250), JUNB (#3753), NFAT1 (#5861), NFAT2 (#8032), NFκB p50 (#12540), and PARP1 (#9532) from Cell Signaling; NFκB p65 (c15310256) from Diagenode; and GAPDH (NB600-502) from Novus were used as primary antibodies. Membranes were washed with TBS-Tween for 5 min 3 times and were incubated with IRDye secondary antibodies in 1/15,000 dilution in blocking buffer for 60 min at RT with shaking. Membranes were washed with TBS-Tween for 5 min 3 times again and with TBS once. Images were taken with an Odyssey Fluorescent Imaging system (LI-COR).

### ATAC sequencing (ATAC-seq)

Fifty thousand naïve or activated T cells were collected and processed in a transposase reaction, and library preparation was as described by Buenrostro *et al*(Buenrostro et al., 2013).

### Chromatin immunoprecipitation (ChIP) and library preparation by ChIPmentation

Fixing solution (50 mM HEPES-KOH [pH 8.0], 100 mM NaCl, 1 mM EDTA, 0.5 mM EGTA, 8.8% formaldehyde) was added directly to T cell culture at a final formaldehyde concentration of 0.88% for fixation. After the incubation for 4 min on ice, 2M glycine solution was added to a final concentration of 125 mM and incubated at room temperature for 10 min to stop fixation. Cells were transferred to tubes and washed with cold PBS twice. All buffers used in the following procedures were supplemented with 1X protease inhibitor solution (SIGMA, cat#P8340) before use. Fixed cells were resuspended in L1 buffer (50 mM HEPES-KOH [pH 8.0], 140 mM NaCl, 1 mM EDTA, 10% glycerol, 0.5% NP-40, 0.25% Triton X-100, 1X protease inhibitors) and incubated on ice for 10 min with shaking every 3 min. Isolated nuclei were incubated with L2 buffer (10 mM Tris-HCl [pH 8.0], 200 mM NaCl, 1 mM EDTA, and 0.5 mM EGTA) with rotation for 10 min at RT. Then, isolated nuclei were carefully washed and resuspended in TE+0.1% SDS solution. Cells were sonicated (peak power: 105.0, Duty Factor: 10.0, Cycle/Burst: 200) for 45 sec in microtubes at 4 °C using S220 Focused ultrasonicators (COVARIS) to obtain the 200-500–bp fragments of chromatin. Chromatin solution was centrifuged, and supernatant chromatin was collected. Triton X-100, glycerol, NaCl, and sodium deoxycholate were added into the chromatin solution to final concentrations of 1%, 5%, 150 mM, and 0.1%, respectively. Protease inhibitors were also added.

ChIPs were performed in an IP-Star® Compact automation system (Diagenode). In brief, about 5-10 μg of chromatin solution corresponding to 1-2 million cells and 5-10 μL of Protein A or G Dynabeads (Thermo) were used per reaction. Antibodies against cFOS (Santa Cruz Biotechnology, sc-7202), H3K27Ac (Diagenode, pAb-196-050), or H3K4me3 (Millipore #17-614) were used. The other antibodies were the same as used in western blotting. The Dynabeads were sequentially washed with wash buffer 1 (RIPA 150 mM NaCl: 10 mM Tris-HCl [pH 8.0], 150 mM NaCl, 1 mM EDTA, 0.1% SDS, 0.1% sodium deoxycholate, and 1% Triton X-100), wash buffer 2 (RIPA 250 mM NaCl), wash buffer 3 (50 mM Tris-HCl [pH 8.0], 2 mM EDTA, and 0.2% N-Lauroylsarcosine sodium salt), and wash buffer 4 (TE+0.2% Triton X-100) for 15 min each. Washed Dynabeads were processed with transposase per ChIPmentation protocol to accomplish tagmentation of ChIPed DNA(Schmidl et al., 2015). Tagmented DNA was incubated with proteinase K in elution buffer (TE with 250 mM NaCl and 0.3% SDS) for 4 h at 65 °C and purified from beads with the Qiagen MinElute DNA kit. PCR amplification and purification were carried out in the same way as for ATAC-seq.

### RNA isolation and RNA sequencing (RNA-seq)

Total RNA was isolated with the Aurum™ Total RNA Mini Kit (BIO-RAD, cat#7326820) including on-column DNaseI digest. PolyA selection and fragmentation of RNA were performed according to the TruSeq^TM^ Stranded polyA RNA Sample Preparation Guide from Illumina. The fragmented RNA was subjected to first-strand cDNA synthesis. For second-strand cDNA synthesis and adapter ligation, the IP-Star® Compact automation system was used. After that, library construction was performed according to the Illumina protocol.

### Sequencing, read mapping, peak calling, peak comparison, and calculation enrichment

DNA libraries for ATAC-seq, ChIP-seq, and RNA-seq were sequenced on HiSeq2500 (Illumina) at the DNA Sequencing and Genotyping Core at Cincinnati Children’s Hospital Medical Center. Sequencing data were deposited to GEO under GSE116698 accession number. Data analysis was performed in BioWardrobe (Kartashov and Barski, 2015). Briefly, 75-bp reads were aligned to the human genome (hg19) using Bowtie (Langmead et al., 2009) (for ATAC-seq and ChIP-seq) or STAR (Dobin et al., 2013) (for RNA-seq). STAR was supplied with RefSeq annotation. For peak calling in ATAC-seq and ChIP-seq, MACS2 (Zhang et al., 2008) software was used.

*Browser screenshots* show the coverage by estimated fragments: reads were extended to estimated (by MACS2) fragment length, normalized to total mapped read number, and displayed as coverage on a mirror of the University of California Santa Cruz (UCSC) genome browser.

*Peak comparison*: MAnorm analysis was performed in BioWardrobe using peaks identified with MACS2 (Shao et al., 2012). To define NORs and EORs, differential fold enrichment > 2 and FDR ≤ 0.05 thresholds were used. For identifying “common” open chromatin regions (CORs), merged common regions from MAnorm analysis with FDR > 0.05 were used.

*Box plots and tag density*: Uniquely mapped reads in ATAC-seq and ChIP-seq samples were extended to MACS2 estimated fragment length. At each nucleotide position, the number of reads mapping there per million of total uniquely mapped reads was calculated and converted into average density over the peak. The density was used for box plots, heatmaps, density profiles, and scatter plots.

*Differential expression analysis*: Differential expression in RNA-seq between samples was identified by DEseq2 software, and the p-value was calculated (Love et al., 2014).

### Motif enrichment analysis

Isolated genomic coordinates of open chromatin regions were used for identifying TF motifs by HOMER software (Heinz et al., 2010). We ran HOMER with the empirical data sets together with the control data sets, which comprised the same length of DNA sequence from the flanking regions within 20,000 bp from each region.

### Gene ontology enrichment analysis and GSEA

Isolated genes nearby open chromatin regions were processed with the ToppFun function in the ToppGene Suite (Chen et al., 2007) website (https://toppgene.cchmc.org/enrichment.jsp) for ontology enrichment analysis. GSEA analysis was performed with manually defined gene sets located next to the open chromatin regions (Mootha et al., 2003; Subramanian et al., 2005).

### Identifying SEs

For identifying SEs, ChIP-seq of H3K27Ac and the same criteria and software as described in Whyte *et al*. and Lovén *et al*. were used (Whyte et al., 2013; Selective Inhibition of Tumor Oncogenes by Disruption of Super-Enhancers, 2013).

### Regulatory element locus intersection (RELI)

Briefly, SNPs that are in linkage disequilibrium with index risk SNPs for various disease from NIH GWAS catalog were identified. RELI software was used to calculate their overlap with ChIP-seq and ATAC-seq peaks with risk loci.

### Analysis of H3K4me2 data in asthmatic patient

H3K4me2 data in Th2 cells from asthmatic donors and healthy controls were collected from GSE53646 (Seumois et al., 2014), and initially processed in BioWardrobe as described above. H3K4me2 data are shown in WUSTL genome browser within BioWardrobe (Zhou et al., 2011). The H3K4me2 differential enrichment domains between asthmatics and healthy controls were identified by using rgt-THOR software (Allhoff et al., 2016).

### A-FOS and GFP protein preparation

A-FOS and GFP cDNA fragments were amplified from CMV500 A-FOS (Addgene, # 33353) and pmaxGFP (Lonza) vectors by PCR, and a 6xHis-GST DNA fragment was also amplified from pGEX vector. A-FOS, GFP, and 6xHis-GST were individually mixed with digested plasmid DNA (GE Healthcare Life Sciences) with *NcoI* restriction enzyme in an infusion cloning reaction (TakaraBio). The constructed plasmid was transformed into the BL21 bacterial strain. A single colony was picked and transferred to 10 mL of TB-SB medium (TB:SB = 1:1) with Kanamycin, cultured overnight at 37 °C, transferred to 500 mL of TB-SB medium, and cultured until the optical density at 600 nm reached 0.8. For protein expression, IPTG was added into the culture to a final concentration of 0.5 mM, and bacteria were cultured for another 6-8 h at 34 °C. The bacteria were spun down, washed with PBS twice, and stored at −80 °C. The bacterial pellet was resuspended in 20 mL lysis buffer (50 mM Na_2_HPO_4_ pH 8, 300 mM NaCl, 10% glycerol, 0.1% Triton X100, 20 mM imidazole with addition of protease inhibitors, lysozyme [100 μg/mL], DNaseI [0.2 μ/mL], and beta-mercaptoethanol to 1 mM), sonicated with a probe sonicator, and then frozen, thawed, and sonicated again. The lysate was cleared by centrifugation and loaded on a 5-mL HisTrap HP column in Buffer A (50 mM Na_2_HPO_4_ pH 8, 300 mM NaCl, 20 mM imidazole) using AKTA Start (GE Healthcare Life Sciences). The bound proteins were eluted by the gradient of Buffer B (50 mM Na_2_HPO_4_ pH 8, 300 mM NaCl, 500 mM imidazole). The fractions containing protein were combined and subjected to dialysis against PBS with 1 mM beta-mercaptoethanol overnight. After dialysis, Triton X100 was added to the protein solution, and the His_6_-GST tag was cut with His-tagged HRV3C protease (Sigma) at RT for 5 h. In order to remove the tags and the protease, imidazole was added to 20 mM, the solution was again passed through a 5-mL HisTrap HP column, and the flow-through was collected. The purity was confirmed by gel electrophoresis (data not shown). The Endotoxin Removal Beads (Miltenyi Biotec cat# 130-093-657) were used to remove endotoxins.

### Protein Electroporation

A-FOS and GFP proteins were electroporated using Neon Nucleofector (Invtirogen) and the Neon™ Transfection System 100 µL Kit (Invitrogen, cat# MPK10025). Isolated naïve cells (3.5 million cells) were mixed with 25 µg of A-FOS or GFP in T buffer and pulsed. For activation, the cells were stimulated 2 h after electroporation as described above.

## Acknowledgements

We thank the Cincinnati Children’s Hospital Medical Center DNA Sequencing core for the next generation sequencing; R. Giulitto, C. Habel, and all blood donors at the University of Cincinnati Hoxworth Blood Center for blood donations; and S. Hottinger for editorial assistance.

## Funding

This research was supported in part by the AAI Careers in Immunology Fellowship (MY); NHLBI Career Transition Award (K22 HL098691, AB); NIH Director’s New Innovator Award (DP2 GM119134, AB); NIH P30 grant DK078392; and NINDS R01 NS099068, Lupus Research Alliance “Novel Approaches”, CCRF Endowed Scholar, CCHMC CpG Pilot study award, and CCHMC Trustee Awards to MTW.

## Author Contributions

M.Y. conducted experiments with support from S.J. and A.B. and performed data analysis with support from A.V.K., X.C., M.T.W., and A.B.; M.Y. and A.B. conceived the project and wrote the paper; and all authors read the paper.

## Declaration of Interests

AK and AB are co-founders of Datirium, LLC.

## Supplemental Figures

**Figure S1.**
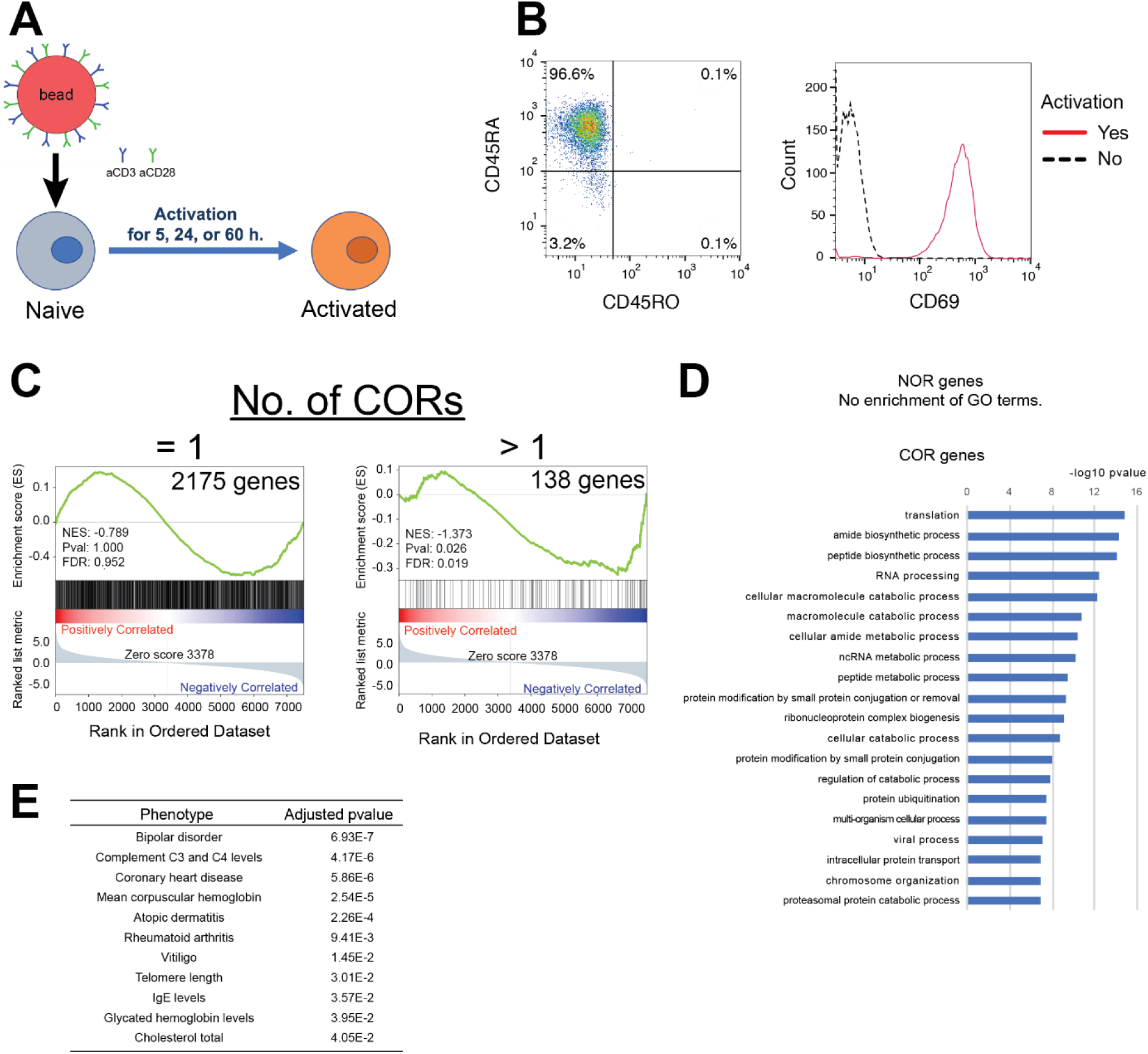
Experimental scheme and characterization of NORs and CORs. (A) Experimental scheme: activation of primary human naive CD4 T cells using beads covered with anti-CD3 and anti-CD28 antibodies. (B) Flow cytometric analysis of isolated human naïve CD4 T cells. Left: counts for CD45RA and CD45RO. Right: counts for CD69 with or without 5 h activation. (C) GSEA analysis compares the gene list ranked by expression fold change during activation with the sets of genes that are located next to 1 or more than 1 COR. NES: Normalized Enrichment Score. (D) Gene ontology analysis of genes adjacent to NORs or CORs. Top gene ontology biological processes (GO) terms and −log10 p-values are shown. (E) Overlap between disease risk SNPs and CORs. Significance of overlap between disease risk SNPs and CORs as calculated by RELI approach.

**Figure S2.**
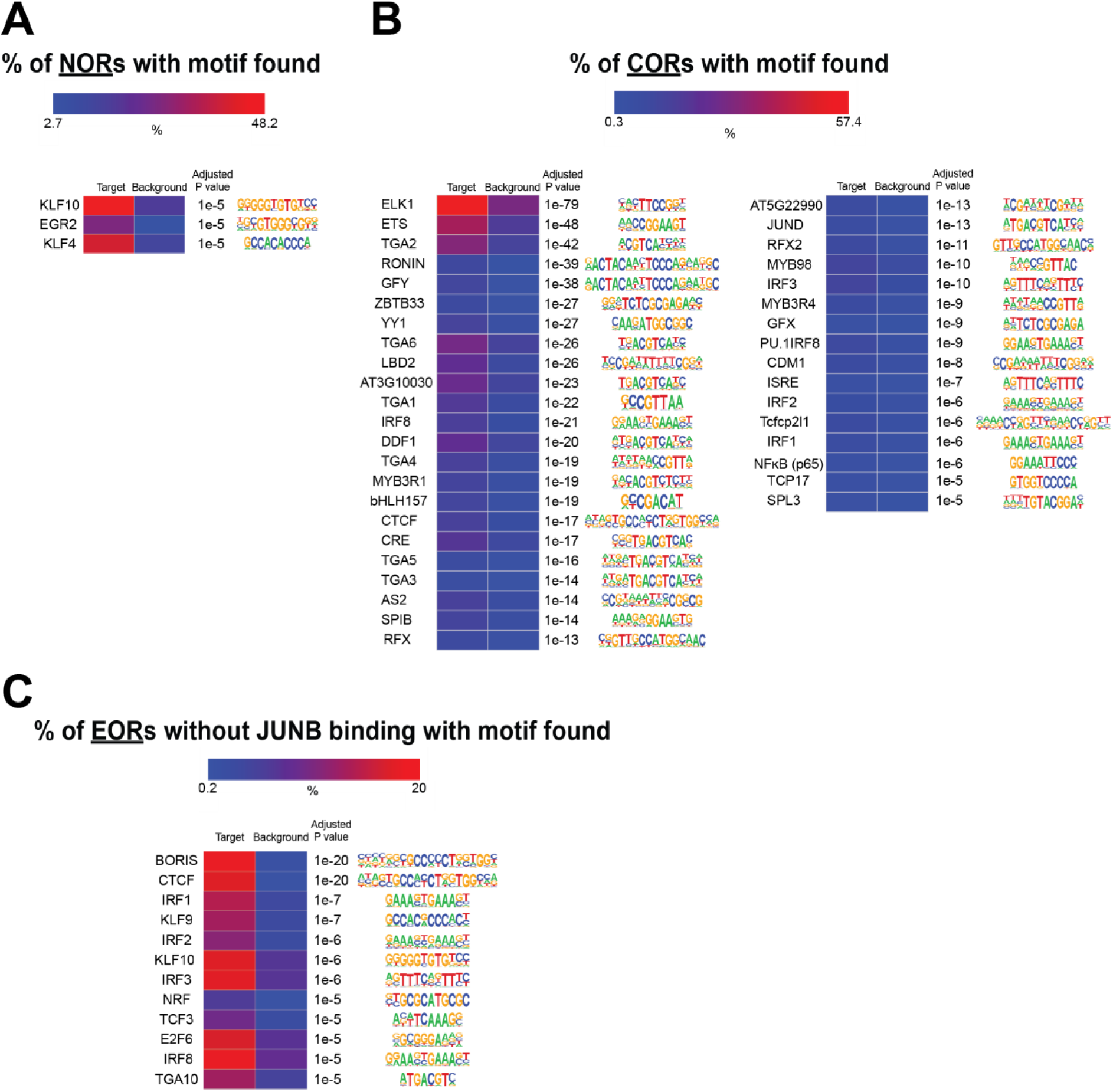
DNA binding motifs enriched in open chromatin regions. (A and B) Enriched DNA motifs in NORs and CORs. Heatmap shows the percentage of regions with motifs. Overrepresented motifs were identified by HOMER analysis and selected with an adjusted p-value ≤ 10^−5^ and target / background > 2. (C) Same analysis as in a and b for EORs *not* bound by JUNB.

**Figure S3.**
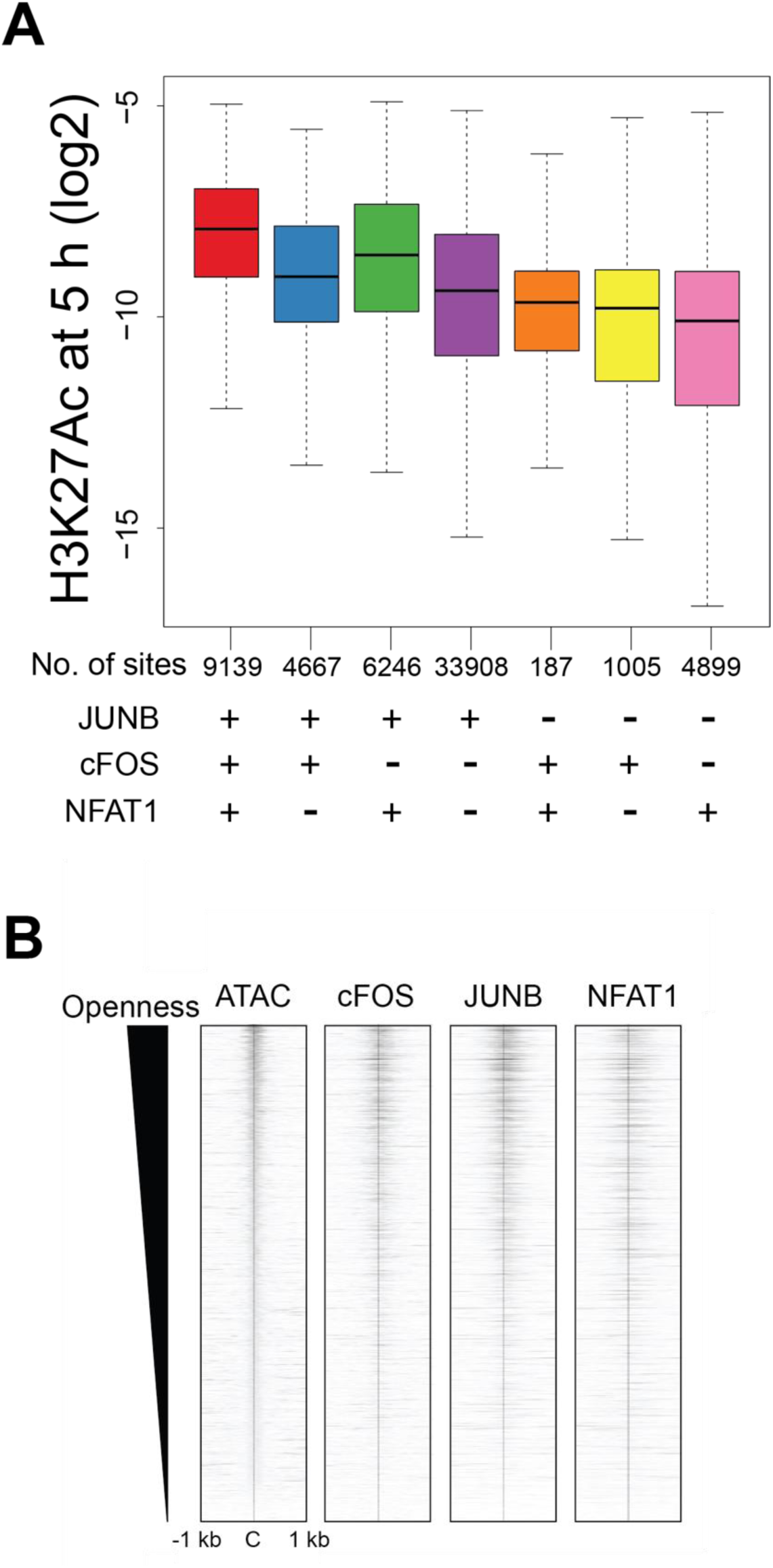
Combinatorial effects of AP-1 and NFAT1 to H3K27Ac and openness. (A) Boxplot showing H3K27Ac ChIP-seq signal in activated cells at 5 h in the regions bound by a given combination of TFs. The y axis shows the H3K27Ac ChIP-seq tag density for TF ChIP-seq peaks. (B) Correlation of openness with TF enrichments in EORs. Heatmap shows the normalized fragment density of ATAC-seq and TF ChIP-seq in a 1-kb radius of the center of the EOR. C: center of open regions.

**Figure S4.**
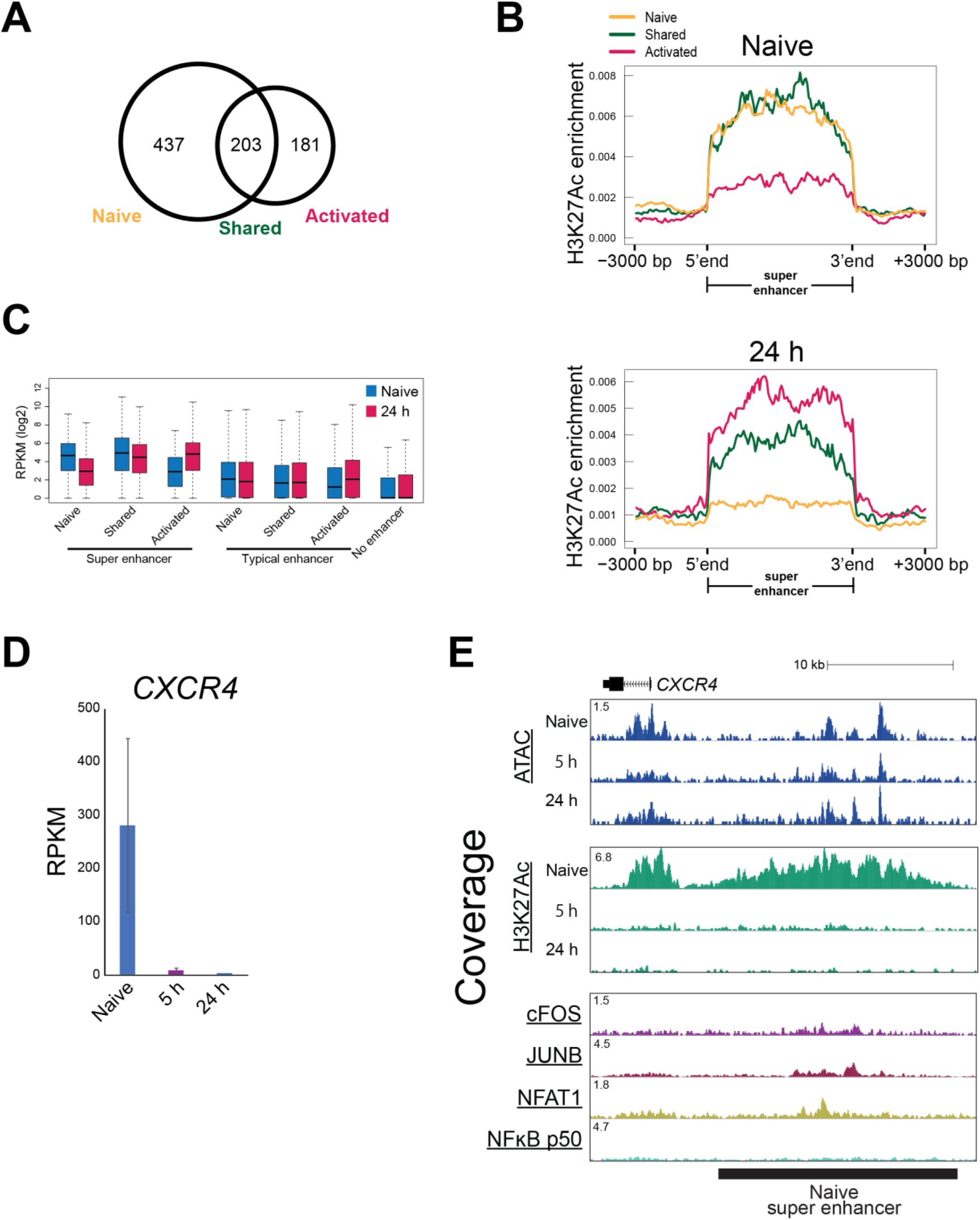
Detection and characterization of super enhancers in naïve and activated T cells. (A) Venn diagram showing the overlap of super enhancers (SEs) in naive and 24-h activated T cells. (B) Average tag density profile of H3K27Ac within SE regions in naive and activated T cells at 24 h. Naive: naive SE, Shared: shared SE, and Activated: activated SE. (C) Boxplot shows expression of genes with naive (Naive), shared (Shared), or activated (Activated) SEs or without SE (No) in naive and 24-h activated T cells. (D) Bar plots show expression of *CXCR4* by RNA-seq. Mean and standard error are shown. n=2. (E) UCSC genome browser screenshot shows ATAC signal and H3K27ac and TFs ChIP-seq at a naive SE in the *CXCR4* locus.

**Figure S5.**
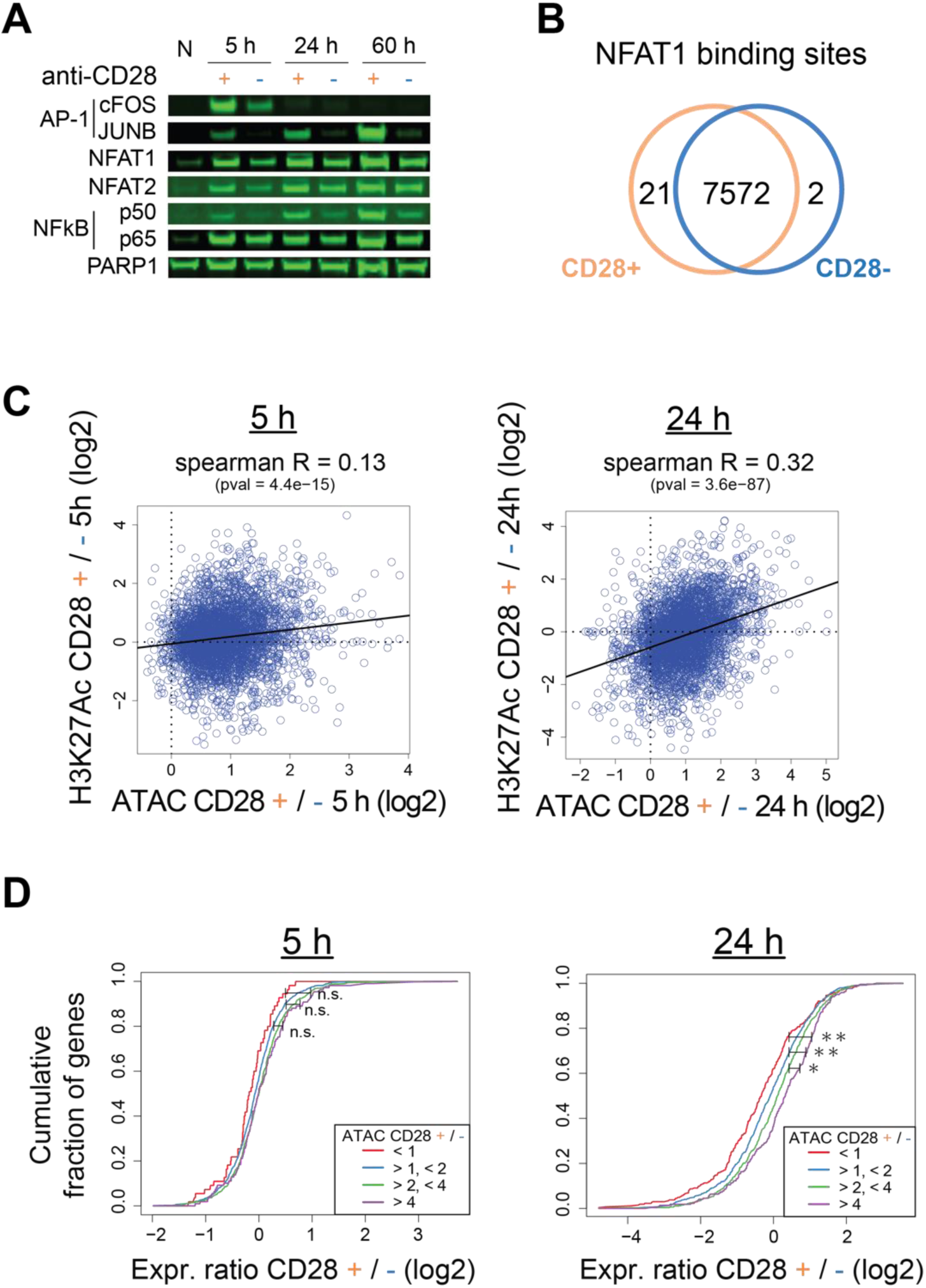
Expression and chromatin changes during anergy induction in the absence of co-stimulation. (A) Western blot shows the nuclear levels of TFs during T cell activation with or without co-stimulation. N: naive T cells. (B) Venn diagram shows the overlap between NFAT1 sites detected in cells stimulated with or without co-stimulation. (B) Correlation between the change in chromatin opening (ATAC) and H3K27Ac (ChIP-seq) at EOR at 5 h (left) and 24 h (right). (G) Cumulative distribution of fold changes between effector (CD28+) and anergic (CD28−) conditions for genes that have indicated ratios of ATAC value in EORs between activated and anergic T cells at 5 h (left) and 24 h (right). X axis: the ratio of expression between E and A conditions (log2). Y axis: the cumulative fraction of genes. Each line indicates gene groups by the ratio of ATAC value (for all EOR peaks that are nearby) (E/A), red < 1, blue > 1 and < 2, green > 2 and < 4, and purple > 4. *p < 0.05 and **p < 0.01 (Kolmogorov–Smirnov test).

**Table S1. List of significant GWAS datasets overlapping NORs, CORs, or EORs.**

## References

Allhoff, M., K. Seré, J. F Pires, M. Zenke, and I. G Costa. 2016. Differential peak calling of ChIP-seq signals with replicates with THOR. Nucleic Acids Research. 15:gkw680–14. doi:10.1093/nar/gkw680.

Armstrong, E.J. 2004. Heart Valve Development: Endothelial Cell Signaling and Differentiation. Circulation Research. 95:459–470. doi:10.1161/01.RES.0000141146.95728.da.

Barski, A., R. Jothi, S. Cuddapah, K. Cui, T.Y. Roh, D.E. Schones, and K. Zhao. 2009. Chromatin poises miRNA- and protein-coding genes for expression. Genome Research. 19:1742–1751. doi:10.1101/gr.090951.109.

Ben Youngblood, J.S. Hale, H.T. Kissick, E. Ahn, X. Xu, A. Wieland, K. Araki, E.E. West, H.E. Ghoneim, Y. Fan, P. Dogra, C.W. Davis, B.T. Konieczny, R. Antia, X. Cheng, and R. Ahmed. 2017. Effector CD8 T cells dedifferentiate into long-lived memory cells. Nature. 552:404–409. doi:10.1038/nature25144.

Bevington, S.L., P. Cauchy, J. Piper, E. Bertrand, N. Lalli, R.C. Jarvis, L.N. Gilding, S. Ott, C. Bonifer, and P.N. Cockerill. 2016. Inducible chromatin priming is associated with the establishment of immunological memory in T cells. EMBO J. 1–21. doi:10.15252/embj.201592534.

Biddie, S.C., S. John, P.J. Sabo, R.E. Thurman, T.A. Johnson, R.L. Schiltz, T.B. Miranda, M.-H. Sung, S. Trump, S.L. Lightman, C. Vinson, J.A. Stamatoyannopoulos, and G.L. Hager. 2011. Transcription Factor AP1 Potentiates Chromatin Accessibility and Glucocorticoid Receptor Binding. Molecular Cell. 43:145–155. doi:10.1016/j.molcel.2011.06.016.

Buenrostro, J.D., P.G. Giresi, L.C. Zaba, H.Y. Chang, and W.J. Greenleaf. 2013. Transposition of native chromatin for fast and sensitive epigenomic profiling of open chromatin, DNA-binding proteins and nucleosome position. Nat. Methods. 10:1213–1218. doi:10.1038/nmeth.2688.

Chen, J., H. Xu, B.J. Aronow, and A.G. Jegga. 2007. Improved human disease candidate gene prioritization using mouse phenotype. BMC Bioinformatics. 8:392–13. doi:10.1186/1471-2105-8-392.

Chen, L., J.N. Glover, P.G. Hogan, A. Rao, and S.C. Harrison. 1998. Structure of the DNA-binding domains from NFAT, Fos and Jun bound specifically to DNA. Nature. 392:42–48. doi:10.1038/32100.

Chou, C., A.K. Pinto, J.D. Curtis, S.P. Persaud, M. Cella, C.-C. Lin, B.T. Edelson, P.M. Allen, M. Colonna, E.L. Pearce, M.S. Diamond, and T. Egawa. 2014. c-Myc-induced transcription factor AP4 is required for host protection mediated by CD8+ T cells. Nature Immunology. 15:884–893. doi:10.1038/ni.2943.

Crabtree, G.R., and E.N. Olson. 2002. NFAT signaling: choreographing the social lives of cells. Cell. 109 Suppl:S67–79.

Djuretic, I.M., D. Levanon, V. Negreanu, Y. Groner, A. Rao, and K.M. Ansel. 2006. Transcription factors T-bet and Runx3 cooperate to activate Ifng and silence Il4 in T helper type 1 cells. Nature Immunology. 8:145–153. doi:10.1038/ni1424.

Dobin, A., C.A. Davis, F. Schlesinger, J. Drenkow, C. Zaleski, S. Jha, P. Batut, M. Chaisson, and T.R. Gingeras. 2013. STAR: ultrafast universal RNA-seq aligner. Bioinformatics. 29:15–21. doi:10.1093/bioinformatics/bts635.

Durant, L., W.T. Watford, H.L. Ramos, A. Laurence, G. Vahedi, L. Wei, H. Takahashi, H.-W. Sun, Y. Kanno, F. Powrie, and J.J.O. Shea. 2010. Diverse Targets of the Transcription Factor STAT3 Contribute to T Cell Pathogenicity and Homeostasis. Immunity. 32:605–615. doi:10.1016/j.immuni.2010.05.003.

Ernst, J., P. Kheradpour, T.S. Mikkelsen, N. Shoresh, L.D. Ward, C.B. Epstein, X. Zhang, L. Wang, R. Issner, M. Coyne, M. Ku, T. Durham, M. Kellis, and B.E. Bernstein. 2011. Mapping and analysis of chromatin state dynamics in nine human cell types. Nature. 473:43–49. doi:10.1038/nature09906.

Farh, K.K.-H., A. Marson, J. Zhu, M. Kleinewietfeld, W.J. Housley, S. Beik, N. Shoresh, H. Whitton, R.J.H. Ryan, A.A. Shishkin, M. Hatan, M.J. Carrasco-Alfonso, D. Mayer, C.J. Luckey, N.A. Patsopoulos, P.L. De Jager, V.K. Kuchroo, C.B. Epstein, M.J. Daly, D.A. Hafler, and B.E. Bernstein. 2014. Genetic and epigenetic fine mapping of causal autoimmune disease variants. Nature. 1–21. doi:10.1038/nature13835.

Fathman, C.G., and N.B. Lineberry. 2007. Molecular mechanisms of CD4+ T-cell anergy. Nat Rev Immunol. 7:599–609. doi:10.1038/nri2131.

Fleischmann, A., F. Hafezi, C. Elliott, C.E. Remé, U. Rüther, and E.F. Wagner. 2000. Fra-1 replaces c-Fos-dependent functions in mice. Genes & Development. 14:2695–2700.

Gruda, M.C., J. van Amsterdam, C.A. Rizzo, S.K. Durham, S. Lira, and R. Bravo. 1996. Expression of FosB during mouse development: normal development of FosB knockout mice. Oncogene. 12:2177–2185.

Harley, J.B., X. Chen, M. Pujato, D. Miller, A. Maddox, C. Forney, A.F. Magnusen, A. Lynch, K. Chetal, M. Yukawa, A. Barski, N. Salomonis, K.M. Kaufman, L.C. Kottyan, and M.T. Weirauch. 2018. Transcription factors operate across disease loci, with EBNA2 implicated in autoimmunity. Nat Genet. 50:699–707. doi:10.1038/s41588-018-0102-3.

Hawkins, R.D., A. Larjo, S.K. Tripathi, U. Wagner, Y. Luu, T. Lönnberg, S.K. Raghav, L.K. Lee, R. Lund, B. Ren, H. Lähdesmäki, and R. Lahesmaa. 2013. Global Chromatin State Analysis Reveals Lineage-Specific Enhancers during the Initiationof Human T helper 1 and T helper 2 Cell Polarization. Immunity. 38:1271–1284. doi:10.1016/j.immuni.2013.05.011.

Heinz, S., C. Benner, N. Spann, E. Bertolino, Y.C. Lin, P. Laslo, J.X. Cheng, C. Murre, H. Singh, and C.K. Glass. 2010. Simple Combinations of Lineage-Determining Transcription Factors Prime cis-Regulatory Elements Required for Macrophage and B Cell Identities. Molecular Cell. 38:576–589. doi:10.1016/j.molcel.2010.05.004.

Ito, T., M. Yamauchi, M. Nishina, N. Yamamichi, T. Mizutani, M. Ui, M. Murakami, and H. Iba. 2001. Identification of SWI{middle dot}SNF Complex Subunit BAF60a as a Determinant of the Transactivation Potential of Fos/Jun Dimers. J. Biol. Chem. 276:2852–2857. doi:10.1074/jbc.M009633200.

Jain, J., E.A. Nalefski, P.G. McCaffrey, R.S. Johnson, B.M. Spiegelman, V. Papaioannou, and A. Rao. 1994. Normal peripheral T-cell function in c-Fos-deficient mice. Molecular and Cellular Biology. 14:1566–1574.

Jauliac, S., C. López-Rodriguez, L.M. Shaw, L.F. Brown, A. Rao, and A. Toker. 2002. The role of NFAT transcription factors in integrin-mediated carcinoma invasion. Nat Cell Biol. 4:540–544. doi:10.1038/ncb816.

Kanhere, A., A. Hertweck, U. Bhatia, M.R.G.O. kmen, E. Perucha, I. Jackson, G.M. Lord, and R.G. Jenner. 1AD. T-bet and GATA3 orchestrate Th1 and Th2 differentiation through lineage-specific targeting of distal regulatory elements. Nature Communications. 3:1–12. doi:10.1038/ncomms2260.

Kartashov, A.V., and A. Barski. 2015. BioWardrobe: an integrated platform for analysis of epigenomics and transcriptomics data. Genome Biol. 16:158. doi:10.1186/s13059-015-0720-3.

Komori, H.K., T. Hart, S.A. LaMere, P.V. Chew, and D.R. Salomon. 2015. Defining CD4 T Cell Memory by the Epigenetic Landscape of CpG DNA Methylation. J. Immunol. 194:1565–1579. doi:10.4049/jimmunol.1401162.

Kriegel, M.A., C. Rathinam, and R.A. Flavell. 2009. E3 ubiquitin ligase GRAIL controls primary T cell activation and oral tolerance. Proc. Natl. Acad. Sci. U.S.A. 106:16770–16775. doi:10.1073/pnas.0908957106.

Kuo, C.T., M.L. Veselits, and J.M. Leiden. 1997. LKLF: A transcriptional regulator of single-positive T cell quiescence and survival. Science. 277:1986–1990. doi:10.1126/science.277.5334.1986.

Langmead, B., C. Trapnell, M. Pop, and S.L. Salzberg. 2009. Ultrafast and memory-efficient alignment of short DNA sequences to the human genome. Genome Biol. 10:R25. doi:10.1186/gb-2009-10-3-r25.

Lee, M.D., K.N. Bingham, T.Y. Mitchell, J.L. Meredith, and J.S. Rawlings. 2015. Calcium mobilization is both required and sufficient for initiating chromatin decondensation during activation of peripheral T-cells. Molecular Immunology. 63:540–549. doi:10.1016/j.molimm.2014.10.015.

Liu, X., C.T. Berry, G. Ruthel, J.J. Madara, K. MacGillivray, C.M. Gray, L.A. Madge, K.A. McCorkell, D.P. Beiting, U. Hershberg, M.J. May, and B.D. Freedman. 2016. T Cell Receptor-induced Nuclear Factor κB (NF-κB) Signaling and Transcriptional Activation Are Regulated by STIM1- and Orai1-mediated Calcium Entry. Journal of Biological Chemistry. 291:8440–8452. doi:10.1074/jbc.M115.713008.

Love, M.I., W. Huber, and S. Anders. 2014. Moderated estimation of fold change and dispersion for RNA-seq data with DESeq2. Genome Biol. 15:31–21. doi:10.1186/s13059-014-0550-8.

Macian, F., C. López-Rodríguez, and A. Rao. 2001. Partners in transcription: NFAT and AP-1. Oncogene. 20:2476–2489. doi:10.1038/sj.onc.1204386.

Macian, F., F. García-Cózar, S.-H. Im, H.F. Horton, M.C. Byrne, and A. Rao. 2002. Transcriptional mechanisms underlying lymphocyte tolerance. Cell. 109:719–731.

Mazzoni, A., V. Santarlasci, L. Maggi, M. Capone, M.C. Rossi, V. Querci, R. De Palma, H.D. Chang, A. Thiel, R. Cimaz, F. Liotta, L. Cosmi, E. Maggi, A. Radbruch, S. Romagnani, J. Dong, and F. Annunziato. 2015. Demethylation of the RORC2 and IL17A in Human CD4+ T Lymphocytes Defines Th17 Origin of Nonclassic Th1 Cells. J. Immunol. 194:3116–3126. doi:10.4049/jimmunol.1401303.

Mootha, V.K., C.M. Lindgren, K.-F. Eriksson, A. Subramanian, S. Sihag, J. Lehar, P. Puigserver, E. Carlsson, M. Ridderstråle, E. Laurila, N. Houstis, M.J. Daly, N. Patterson, J.P. Mesirov, T.R. Golub, P. Tamayo, B. Spiegelman, E.S. Lander, J.N. Hirschhorn, D. Altshuler, and L.C. Groop. 2003. PGC-1alpha-responsive genes involved in oxidative phosphorylation are coordinately downregulated in human diabetes. Nat Genet. 34:267–273. doi:10.1038/ng1180.

Mukasa, R., A. Balasubramani, Y.K. Lee, S.K. Whitley, B.T. Weaver, Y. Shibata, G.E. Crawford, R.D. Hatton, and C.T. Weaver. 2010. Epigenetic instability of cytokine and transcription factor gene loci underlies plasticity of the T helper 17 cell lineage. Immunity. 32:616–627. doi:10.1016/j.immuni.2010.04.016.

Muthusamy, N., K. Barton, and J.M. Leiden. 1995. Defective activation and survival of T cells lacking the Ets-1 transcription factor. Nature. 377:639–642. doi:10.1038/377639a0.

Nguyen, T.N., L.J. Kim, R.D. Walters, L.F. Drullinger, T.N. Lively, J.F. Kugel, and J.A. Goodrich. 2010. The C-terminal region of human NFATc2 binds cJun to synergistically activate interleukin-2 transcription. Molecular Immunology. 47:2314–2322. doi:10.1016/j.molimm.2010.05.287.

Ohkura, N., M. Hamaguchi, H. Morikawa, K. Sugimura, A. Tanaka, Y. Ito, M. Osaki, Y. Tanaka, R. Yamashita, N. Nakano, J. Huehn, H.J. Fehling, T. Sparwasser, K. Nakai, and S. Sakaguchi. 2012. T cell receptor stimulation-induced epigenetic changes and Foxp3 expression are independent and complementary events required for Treg cell development. Immunity. 37:785–799. doi:10.1016/j.immuni.2012.09.010.

Ono, M., H. Yaguchi, N. Ohkura, I. Kitabayashi, Y. Nagamura, T. Nomura, Y. Miyachi, T. Tsukada, and S. Sakaguchi. 2007. Foxp3 controls regulatory T-cell function by interacting with AML1/Runx1. Nature. 446:685–689. doi:10.1038/nature05673.

O’Shea, J.J., and W.E. Paul. 2010. Mechanisms underlying lineage commitment and plasticity of helper CD4+ T cells. Science. 327:1098–1102. doi:10.1126/science.1178334.

Panagoulias, I., T. Georgakopoulos, I. Aggeletopoulou, M. Agelopoulos, D. Thanos, and A. Mouzaki. 2016. Transcription Factor Ets-2 Acts as a Preinduction Repressor of Interleukin-2 (IL-2) Transcription in Naive T Helper Lymphocytes. J. Biol. Chem. 291:26707–26721. doi:10.1074/jbc.M116.762179.

Pham, D., Q. Yu, C.C. Walline, R. Muthukrishnan, J.S. Blum, and M.H. Kaplan. 2013. Opposing Roles of STAT4 and Dnmt3a in Th1 Gene Regulation. J. Immunol. 191:902–911. doi:10.4049/jimmunol.1203229.

Ranger, A.M., L.C. Gerstenfeld, J. Wang, T. Kon, H. Bae, E.M. Gravallese, M.J. Glimcher, and L.H. Glimcher. 2000. The nuclear factor of activated T cells (NFAT) transcription factor NFATp (NFATc2) is a repressor of chondrogenesis. J. Exp. Med. 191:9–22.

Rawlings, J.S., M. Gatzka, P.G. Thomas, and J.N. Ihle. 2010. Chromatin condensation via the condensin II complex is required for peripheral T-cell quiescence. EMBO J. 30:263–276. doi:10.1038/emboj.2010.314.

Rochman, Y., M. Yukawa, A.V. Kartashov, and A. Barski. 2015. Functional Characterization of Human T Cell Hyporesponsiveness Induced by CTLA4-Ig. PLoS ONE. 10:e0122198–18. doi:10.1371/journal.pone.0122198.

Rooney, J.W., Y.L. Sun, L.H. Glimcher, and T. Hoey. 1995. Novel NFAT sites that mediate activation of the interleukin-2 promoter in response to T-cell receptor stimulation. Molecular and Cellular Biology. 15:6299–6310.

Russ, B.E., J.E. Prier, S. Rao, and S.J. Turner. 2013. T cell immunity as a tool for studying epigenetic regulation of cellular differentiation. Front Genet. 4:218. doi:10.3389/fgene.2013.00218.

Safford, M., S. Collins, M.A. Lutz, A. Allen, C.-T. Huang, J. Kowalski, A. Blackford, M.R. Horton, C. Drake, R.H. Schwartz, and J.D. Powell. 2005. Egr-2 and Egr-3 are negative regulators of T cell activation. Nature Immunology. 6:472–480. doi:10.1038/ni1193.

Sareneva, T., S. Matikainen, J. Vanhatalo, K. Melén, J. Pelkonen, and I. Julkunen. 1998. Kinetics of cytokine and NFAT gene expression in human interleukin-2-dependent T lymphoblasts stimulated via T-cell receptor. Immunology. 93:350–357.

Schmidl, C., A.F. Rendeiro, N.C. Sheffield, and C. Bock. 2015. ChIPmentation: fast, robust, low-input ChIP-seq for histones and transcription factors. Nat. Methods. 12:963–965. doi:10.1038/nmeth.3542.

Sekimata, M., M. PErez-Melgosa, S.A. Miller, A.S. Weinmann, P.J. Sabo, R. Sandstrom, M.O. Dorschner, J.A. Stamatoyannopoulos, and C.B. Wilson. 2009. CCCTC-Binding Factor and the Transcription Factor T-bet Orchestrate T Helper 1 Cell-SpecificStructure and Function at the Interferon-g Locus. Immunity. 31:551–564. doi:10.1016/j.immuni.2009.08.021.

Selective Inhibition of Tumor Oncogenes by Disruption of Super-Enhancers. 2013. Selective Inhibition of Tumor Oncogenes by Disruption of Super-Enhancers. 153:320–334. doi:10.1016/j.cell.2013.03.036.

Seumois, G., L. Chavez, A. Gerasimova, M. Lienhard, N. Omran, L. Kalinke, M. Vedanayagam, A.P.V. Ganesan, A. Chawla, R. Djukanović, K.M. Ansel, B. Peters, A. Rao, and P. Vijayanand. 2014. Epigenomic analysis of primary human T cells reveals enhancers associated with TH2 memory cell differentiation and asthma susceptibility. Nature Immunology. 15:777–788. doi:10.1038/ni.2937.

Shao, Z., Y. Zhang, G.-C. Yuan, S.H. Orkin, and D.J. Waxman. 2012. MAnorm: a robust model for quantitative comparison of ChIP-Seq data sets. Genome Biol. 13:R16. doi:10.1186/gb-2012-13-3-r16.

Skerka, C., E.L. Decker, and P.F. Zipfel. 1995. A regulatory element in the human interleukin 2 gene promoter is a binding site for the zinc finger proteins Sp1 and EGR-1. J. Biol. Chem. 270:22500–22506.

Smith, A.E., C. Chronis, M. Christodoulakis, S.J. Orr, N.C. Lea, N.A. Twine, A. Bhinge, G.J. Mufti, and N.S.B. Thomas. 2009. Epigenetics of human T cells during the G0->G1 transition. Genome Research. 19:1325–1337. doi:10.1101/gr.085530.108.

Soderquest, K., A. Hertweck, C. Giambartolomei, S. Henderson, R. Mohamed, R. Goldberg, E. Perucha, L. Franke, J. Herrero, V. Plagnol, R.G. Jenner, and G.M. Lord. 2017. Genetic variants alter T-bet binding and gene expression in mucosal inflammatory disease. PLoS Genetics. 13:e1006587–23. doi:10.1371/journal.pgen.1006587.

Subramanian, A., P. Tamayo, V.K. Mootha, S. Mukherjee, B.L. Ebert, M.A. Gillette, A. Paulovich, S.L. Pomeroy, T.R. Golub, E.S. Lander, and J.P. Mesirov. 2005. Gene set enrichment analysis: a knowledge-based approach for interpreting genome-wide expression profiles. Proceedings of the National Academy of Sciences. 102:15545– 15550. doi:10.1073/pnas.0506580102.

Thaker, Y.R., H. Schneider, and C.E. Rudd. 2015. TCR and CD28 activate the transcription factor NF-κB in T-cells via distinct adaptor signaling complexes. Immunology Letters. 163:113–119. doi:10.1016/j.imlet.2014.10.020.

Trushin, S.A., K.N. Pennington, A. Algeciras-Schimnich, and C.V. Paya. 1999. Protein kinase C and calcineurin synergize to activate IkappaB kinase and NF-kappaB in T lymphocytes. J. Biol. Chem. 274:22923–22931.

Turner, H., M. Gomez, E. McKenzie, A. Kirchem, A. Lennard, and D.A. Cantrell. 1998. Rac-1 regulates nuclear factor of activated T cells (NFAT) C1 nuclear translocation in response to Fcepsilon receptor type 1 stimulation of mast cells. J. Exp. Med. 188:527–537.

Vahedi, G., H. Takahashi, S. Nakayamada, H.-W. Sun, V. Sartorelli, Y. Kanno, and J.J. O’Shea. 2012. STATs Shape the Active Enhancer Landscape of T Cell Populations. Cell. 151:981–993. doi:10.1016/j.cell.2012.09.044.

Vierbuchen, T., E. Ling, C.J. Cowley, C.H. Couch, X. Wang, D.A. Harmin, C.W.M. Roberts, and M.E. Greenberg. 2017. AP-1 Transcription Factors and the BAF Complex Mediate Signal-Dependent Enhancer Selection. Molecular Cell. 68:1067–1082.e12. doi:10.1016/j.molcel.2017.11.026.

Walters, R.D., L.F. Drullinger, J.F. Kugel, and J.A. Goodrich. 2013. NFATc2 recruits cJun homodimers to an NFAT site to synergistically activate interleukin-2 transcription. Molecular Immunology. 56:48–56. doi:10.1016/j.molimm.2013.03.022.

Wan, C.-K., A.B. Andraski, R. Spolski, P. Li, M. Kazemian, J. Oh, L. Samsel, P.A. Swanson II, D.B. McGavern, E.P. Sampaio, A.F. Freeman, J.D. Milner, S.M. Holland, and W.J. Leonard. 2015. Opposing roles of STAT1 and STAT3 in IL-21 function in CD4 +T cells. Proceedings of the National Academy of Sciences. 112:9394–9399. doi:10.1073/pnas.1511711112.

Wang, R., C.P. Dillon, L.Z. Shi, S. Milasta, R. Carter, D. Finkelstein, L.L. McCormick, P. Fitzgerald, H. Chi, J. Munger, and D.R. Green. 2011. The Transcription Factor Myc Controls Metabolic Reprogramming upon T Lymphocyte Activation. Immunity. 35:871–882. doi:10.1016/j.immuni.2011.09.021.

Wei, G., L. Wei, J. Zhu, C. Zang, J. Hu-Li, Z. Yao, K. Cui, Y. Kanno, T.-Y. Roh, W.T. Watford, D.E. Schones, W. Peng, H.-W. Sun, W.E. Paul, J.J. O’Shea, and K. Zhao. 2009. Global Mapping of H3K4me3 and H3K27me3 Reveals Specificity and Plasticity in LineageFate Determination of Differentiating CD4. Immunity. 30:155–167. doi:10.1016/j.immuni.2008.12.009.

Whyte, W.A., D.A. Orlando, D. Hnisz, B.J. Abraham, C.Y. Lin, M.H. Kagey, P.B. Rahl, T.I. Lee, and R.A. Young. 2013. Master Transcription Factors and Mediator Establish Super-Enhancers at Key Cell Identity Genes. Cell. 153:307–319. doi:10.1016/j.cell.2013.03.035.

Wong, H.K., G.M. Kammer, G. Dennis, and G.C. Tsokos. 1999. Abnormal NF-kappa B activity in T lymphocytes from patients with systemic lupus erythematosus is associated with decreased p65-RelA protein expression. J. Immunol. 163:1682–1689.

Xu, Y.Z., T. Thuraisingam, R. Marino, and D. Radzioch. 2011. Recruitment of SWI/SNF complex is required for transcriptional activation of the SLC11A1 gene during macrophage differentiation of HL-60 cells. Journal of Biological Chemistry. 286:12839–12849. doi:10.1074/jbc.M110.185637.

Yamada, T., C.S. Park, M. Mamonkin, and H.D. Lacorazza. 2009. Transcription factor ELF4 controls the proliferation and homing of CD8+ T cells via the Krüppel-like factors KLF4 and KLF2. Nature Immunology. 10:618–626. doi:10.1038/ni.1730.

Zaret, K.S., and S.E. Mango. 2016. Pioneer transcription factors, chromatin dynamics, and cell fate control. Current Opinion in Genetics & Development. 37:76–81. doi:10.1016/j.gde.2015.12.003.

Zhang, Y., T. Liu, C.A. Meyer, J. Eeckhoute, D.S. Johnson, B.E. Bernstein, C. Nusbaum, R.M. Myers, M. Brown, W. Li, and X.S. Liu. 2008. Model-based analysis of ChIP-Seq (MACS). Genome Biol. 9:R137. doi:10.1186/gb-2008-9-9-r137.

Zhou, X., B. Maricque, M. Xie, D. Li, V. Sundaram, E.A. Martin, B.C. Koebbe, C. Nielsen, M. Hirst, P. Farnham, R.M. Kuhn, J. Zhu, I. Smirnov, W.J. Kent, D. Haussler, P.A.F. Madden, J.F. Costello, and T. Wang. 2011. The Human Epigenome Browser at Washington University. Nat. Methods. 8:989–990. doi:10.1038/nmeth.1772.

Zhu, J., and W.E. Paul. 2010. Peripheral CD4+ T-cell differentiation regulated by networks of cytokines and transcription factors. Immunological Reviews. 238:247–262. doi:10.1111/j.1600-065X.2010.00951.x.

Zhu, J., H. Yamane, and W.E. Paul. 2010. Differentiation of Effector CD4 T Cell Populations *. Annu. Rev. Immunol. 28:445–489. doi:10.1146/annurev-immunol-030409-101212.

